# Genomic localization bias of secondary metabolite gene clusters and association with histone modifications in *Aspergillus*

**DOI:** 10.1101/2024.02.20.581327

**Authors:** Xin Zhang, Iseult Leahy, Jérȏme Collemare, Michael F. Seidl

**Affiliations:** Theoretical Biology & Bioinformatics Group, Department of Biology, Utrecht University, Padualaan 8, 3584 CH Utrecht, the Netherlands; Westerdijk Fungal Biodiversity Institute, Uppsalalaan 8, 3584 CT Utrecht, the Netherlands

**Keywords:** Aspergilli, secondary metabolite backbone gene, synteny, histone post-transcriptional modification, RNA-seq

## Abstract

Fungi are well-known producers of bioactive secondary metabolites (SMs), which have been exploited for decades by humankind for various medical applications like therapeutics and antibiotics. SMs are synthesized by biosynthetic gene clusters (BGCs) – physically co-localized and co-regulated genes. Because BGCs are often regulated by histone post-translational modifications (PTMs), it was suggested that their chromosomal location is important for their expression. Studies in a few fungal species indicated an enrichment of BGCs in sub-telomeric regions; however, there is no evidence that BGCs with distinct genomic localization are regulated by different histone PTMs. Here, we used 174 *Aspergillus* species covering 22 sections to determine the correlation between BGC genomic localization, gene expression and histone PTMs. We found a high abundance and diversity of SM backbone genes across the *Aspergillus* genus, with notable unique genes within sections. Being unique or conserved in many species, BGCs showed a strong bias for being localized in low-synteny regions, regardless of their position in chromosomes. Using chromosome-level assemblies, we also confirmed a significantly biased localization in sub-telomeric regions. Notably, SM backbone genes in sub-telomeric regions and about half of those in low-synteny regions exhibit higher gene expression variability, likely due to the similar higher variability in H3K4me3 and H3K36me3 histone PTMs; while variations in histone H3 acetylation and H3K9me3 are not correlated to genomic localization and expression variation, as analyzed in two *Aspergillus* species. Expression variability across four *Aspergillus* species further supports that BGCs tend to be located in low-synteny regions and that regulation of expression in those regions likely involves different histone PTMs than the most commonly studied modifications.

**Significance:** Fungi are known for producing an array of bioactive compounds with medical benefits, yet our understanding of how the production of these compounds is regulated remains limited. Here, we focused on the fungal genus *Aspergillus*, containing many species known to be prolific producers of bioactive compounds, to systematically uncover the diversity and genomic localization of biosynthetic pathways. By expanding our knowledge beyond the few commonly studied fungal species, this research offers novel insights into how the genomic localization of biosynthetic pathways matters for the regulation of their expression. Thanks to a new view on BGC localization and expression in relation to histone modifications, our results are expected to stimulate functional research on neglected histone modifications that will support the discovery and harnessing of new fungal metabolites for medical and industrial applications.

## Introduction

Filamentous fungi are prolific producers of bioactive molecules, termed secondary metabolites (SMs) or natural products (Mosunova et al., 2021). While these metabolites are not directly necessary to sustain fungal growth, development or reproduction, they play indispensable roles in inter-organismal interactions and fungal survival in diverse ecological niches (Keller, 2019). SMs have multifaceted functions, from acting as pigments that shield fungal spores from UV radiation, and serving as virulence factors in pathogenic interactions, to providing competitive edges through antibiotic properties (Keller, 2019). Humans have harnessed the bioactive potential of these metabolites to significantly enhance their quality of life (Cairns et al., 2021). Fungal SMs have found significant applications in medical treatments, serving as therapeutic agents, exemplified by lovastatin and ergotamine, as well as immunosuppressants like cyclosporine (Newman & Cragg, 2016; Wiemann & Keller, 2014). They also play crucial roles as antibiotics, with penicillin being the most notable example, and as antifungal agents, such as griseofulvin (Newman & Cragg, 2016).

The biosynthesis of fungal SMs is orchestrated by a series of enzymatic reactions. First, backbone (or core) enzymes use precursors derived from primary metabolism to synthesize an intermediate that is then further modified by so-called tailoring enzymes to produce the final compound(s) (Mosunova et al., 2021). Backbone enzymes responsible for the synthesis of diverse SMs have been classified into categories based on their enzymatic functions such as polyketide synthase (PKS), non-ribosomal peptide synthetase (NRPS), PKS-NRPS hybrid enzymes, dimethylallyl tryptophan synthetase (DMATS), terpene cyclase (TC) (Mosunova et al., 2021), and ribosomally synthesized and post-translationally modified peptides (RiPPs) synthetase (Kessler & Chooi, 2022). Genes encoding backbone and tailoring enzymes, as well as genes responsible for transcription control, transport, and/or self-resistance, are co-regulated and often physically co-localized along the chromosome in biosynthetic gene clusters (BGCs) (Mosunova et al., 2021).

The discrepancy between the predicted bioactive potential based on genome mining and the number of known molecules effectively produced can be explained by the tight transcriptional control of SM biosynthetic pathways (Brakhage, 2013). In addition to the regulation by local and global transcription factors (Chiang et al., 2009; Macheleidt et al., 2016), SM production is regulated globally by chromatin dynamics that, in response to both internal and external stimuli, transitions between different chromatin states: namely the ‘open’ euchromatin, loosely packed and transcriptionally active state, and the ‘closed’ heterochromatin, tightly packed and transcriptionally silent state (Collemare & Seidl, 2019; Gacek & Strauss, 2012). Histone post-translational modifications (PTMs) – corresponding to chemical modifications on specific amino acids localized on the ‘tails’ of histone proteins - contribute to the control of the chromatin status. Most histone acetylations and some histone methylations on H3K4 (histone protein H3 at lysine 4) and H3K36 are associated with euchromatin, while histone methylations on H3K9 and H3K27 are associated with heterochromatin (Lawrence et al., 2016). The deletion of histone deacetylases, which changes the chromatin from ‘closed’ heterochromatin to ‘open’ euchromatin, increases the activity of nine out of 14 NRPSs in *Aspergillus fumigatus* (Lee et al., 2009) and the production of kojic acid in *Aspergillus niger* (Li et al., 2019). The knockout of histone acetyltransferases also leads to defects in SM biosynthesis in *Aspergillus nidulans* (Cánovas et al., 2014). Using inhibitors of histone PTMs like SAHA (suberoylanilide hydroxamic acid, inhibiting histone deacetylases) and 5-aza (5-azacytidine, inhibiting DNA methylation) has been widely employed to deregulate the expression of BGCs in diverse fungi (Brakhage, 2013).

Following the discovery that histone PTMs play an important role in the regulation of SM biosynthesis, it was hypothesized that the chromosomal location of BGCs was crucial for their regulation (Palmer and Keller, 2010). Indeed, several studies reported that BGCs tend to be preferentially located at, or close to, sub-telomeric regions, such as in *Fusarium graminearum* (Connolly et al., 2013), and in some well-studied Aspergilli: namely *Aspergillus oryzae* (Kjærbølling et al., 2020), *Aspergillus flavus* (Yang et al., 2022), *A. fumigatus* (Keller, 2019; Perrin et al., 2007), and *A. nidulans* (Klejnstrup et al., 2012). Sub-telomeric regions in fungi are rich in repetitive sequences such as transposable elements and are typically associated with histone PTMs linked to heterochromatin, especially with the presence of H3K9me3 and H3K27me3 (Keller, 2019; Palmer & Keller, 2010; Wiemann et al., 2013). For instance, in *Fusarium fujikuroi*, the PKS19 BGC, which produces fujikurins, is embedded within a region rich in H3K9me3 on the long arm of chromosome VIII (Wiemann et al., 2009). Similarly, in *A. nidulans*, several BGCs occur in proximity to H3K9me3-rich regions (Gacek-Matthews et al., 2016). Thus, the peculiar genomic localization of BGCs might explain why these BGCs remain transcriptionally silent. Moreover, because of their high repeat content, sub-telomeric regions are prone to rearrangements and typically exhibit low synteny, characterized by reduced conservation of physical co-localization of genes compared to other species (Nakao et al., 2009). In *A. nidulans*, compared to *A. oryzae* and *A. fumigatus*, BGCs often occur in less conserved and heavily rearranged genomic regions, even though the majority (77–79%) of the genomes show conserved synteny (Inglis et al., 2013). However, not all SM gene clusters are associated with heterochromatin and sub-telomeric regions. While previous studies have predominantly focused on a limited number of well-known *Aspergillus* sections and species, often finding an association of SM gene clusters with heterochromatin and sub-telomeric regions, it is crucial to note that not all SM gene clusters are confined to these areas. To determine how the genomic localization and regulation of BGCs are linked, we examined the correlation among the BGC diversity, their genomic organizations, gene expression, and histone PTMs together. We here include 174 diverse Aspergilli species covering 22 sections to provide a more nuanced analysis of the distribution, conservation, and genomic localization of SM genes, particularly in relation to genome-wide syntenic regions and histone PTMs using a sub-set of species with chromosome-level assemblies. This extensive approach allows us to explore the implications of BGC localization and histone PTMs on SM gene expression regulation.

## Results and discussions

### A comprehensive phylogenetic analysis of the fungal genus *Aspergillus*

To provide a comprehensive analysis of BGCs in Aspergilli, we included 174 publicly available Aspergilli genomes (Kjærbølling et al., 2018, 2020; Rasmussen, 2017; Theobald, 2018; Theobald et al., 2018; Vesth et al., 2018) and three *Penicillium* species derived from the Joint Genome Institute (Table S1) as outgroups. The assembled genomes range in size from 23.2 to 42.9 Mb and 90% of them fall within the range of 25.8 to 38.8 Mb (Figure 1, Table S1). Four *Aspergillus* species - *A. fumigatus*, *A. nidulans*, *A. niger*, and *Aspergillus ochraceus* - have chromosomal-level genome assemblies (Nierman et al., 2005), which are comprised of eight chromosomes (Figure 1, Table S1). The other well-studied model system, *A. oryzae*, has also a good genome assembly of eleven scaffolds. Twenty-four Aspergilli assemblies have 100 or fewer scaffolds, while *Aspergillus rambellii* has the most fragmented assembly with 4,177 scaffolds (Figure 1, Table S1). The number of predicted protein-coding genes ranges from 7,761 to 15,687, with 90% of Aspergilli having between 9,561 and 14,061 genes (Figure 1, Table S1). Because the genome contiguity and the number of genes varied between different Aspergilli, we sought to calculate the genome completeness by querying each genome assembly for the occurrence of single-copy orthologous BUSCO genes (Table S2). Importantly, we observed very high genome completeness with 90% of all Aspergilli scoring higher than 98.0% BUSCO completeness (Figure 1, Table S1). Even *A. rambellii* with the most fragmented assembly and the fewest genes (7,761) attains a high genome completeness of 97.1%. Similarly, four other Aspergilli with more than 1,000 scaffolds also have high genome completeness (*Aspergillus udagawae* (97.8%), *Aspergillus avenaceus* (99.1%), *Aspergillus haitiensis* (98.3%), and *Aspergillus coremiiformis* (99.5%)) (Figure 1, Table S1). Although some regions might have been missed in highly fragmented genomes, we considered all species included in this study exhibit sufficient quality to perform accurate comparative analyses.

**Figure 1.**
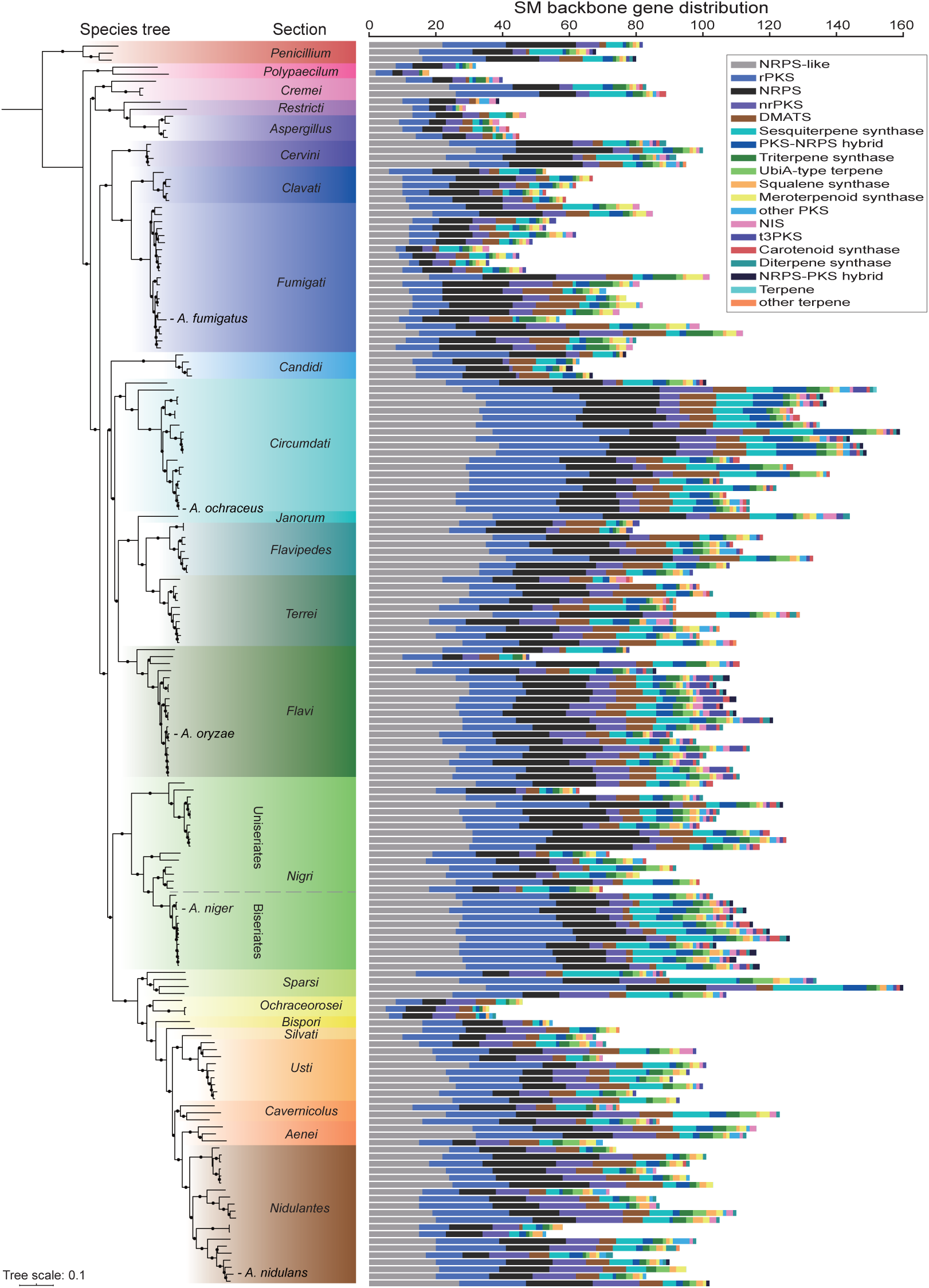
Phylogenetic relationship between *Aspergillus* specie and their secondary metabolite (SM) backbone gene complement. The phylogenetic relationships between 174 *Aspergillus* and three *Penicillium* species are built using a maximum-likelihood framework implemented in IQ-TREE using a concatenated alignment of 758 single-copy BUSCO genes. The dots on the branches indicate ultrafast bootstrap values over 95 and SH-aLRT (Shimodaira-Hasegawa-like approximate likelihood ratio test) values over 80. Species from the same section are highlighted by the same background color. The names of five well-studied Aspergilli are shown in the tree and a dashed line highlights the separation of sub-groups within the *Nigri* section. The smallest and largest dot sizes correspond to the minimum and maximum number in each category, respectively. Genome completeness is indicated by the percentage of identified complete, single-copy BUSCO genes in each genome assembly. SM backbone genes were first detected with antiSMASH, then reclassified by BGCtoolkit. Bar plots representing different SM backbone groups are arranged from left to right from the most to the least abundant.

To provide a robust phylogenetic framework to study BGC and genomic diversity in Aspergilli, we constructed a species phylogeny based on the concatenated alignment of 758 conserved, single-copy BUSCO genes (in total 132,397 informative sites in gene alignments). The tree confirms 22 distinct *Aspergillus* sections, each supported by bootstrap values exceeding 99% (Figure 1), suggesting that our phylogenic analysis is robust. Moreover, this species phylogeny confirms the associations among distinct *Aspergillus* sections as reported in previous studies(Steenwyk et al., 2024). The tree enlarges the current taxonomic breadth as we included a much larger set of diverse 174 *Aspergillus* species compared to less than 90 Aspergilli in previous publications (Steenwyk et al., 2019; Visagie et al., 2014, 2023; Zhang et al., 2022). The 22 sections exhibit notable variation in the number of sequenced species: section *Nigri* has 28 sequenced species, whereas sections *Janorum*, *Bispori*, and *Silvati* are each represented by only a single sequenced species (Figure 1). Despite these differences, the genome assemblies in most sections are relatively uniform in regard to assembly length, number of scaffolds, genome completeness, and gene count, and thus provide a comprehensive and robust reference to study the diversity and evolution of Aspergilli and as well as a solid framework for further analysis.

### Aspergilli harbor an abundant secondary metabolite biosynthetic gene cluster repertoire

To evaluate the potential for SM biosynthesis encoded in the genomes of *Aspergillus* species, we focused on SM backbone genes as these genes are crucial for the first step in SM biosynthesis (Keller, 2019). The focus on SM backbone genes also prevents inaccurate comparison of BGC split over several scaffolds in fragmented assemblies. We used antiSMASH (Blin et al., 2021) to identify regions that comprise a predicted BGC, and employed an in-house BGCtoolkit to further classify SM backbone enzymes into groups that correspond to major chemical families and to retrieve their sequences.

We identified in total 16,243 SM backbone genes across 174 *Aspergillus* species and the three selected *Penicillium*, ranging from 18 in *Aspergillus caninus* to up to 160 in *Aspergillus implicatus* (Figure 1, Table S3). On average, we identified 92 SM backbone genes per Aspergilli compared to only 77 per *Penicillium* species. These SM backbone genes are unevenly distributed. The 41 early-diverging Aspergilli (comprising sections *Polypaecilum*, *Cremei*, *Restricti*, *Aspergillus*, *Cervini*, *Clavati*, and *Fumigati*) have on average 65 SM backbone genes, while the other 133 Aspergilli have on average 99. The *Circumdati* section has the highest average number of SM backbone genes, with its 18 species encoding an average of 129 SM backbone genes. As one BGC can contain several SM backbone genes, the average number of SM backbone genes in Aspergilli is higher than the number of predicted BGCs that have been reported in previous studies. For instance, studies that focused on sections *Nigri* (Theobald et al., 2018), *Flavi* (Kjærbølling et al., 2020) using a customized SMURF approach have detected on average 82 and 70 BGCs, respectively. Here, we found these sections have an average of 104 (section *Nigri*) and 101 (section *Flavi*) SM backbone genes detected with antiSMASH. The tool for BGC prediction greatly influences the numbers as an average of 83 SM backbone genes were reported in section *Terrei* with the SMURF-derived prediction(Theobald et al., 2024), while we predicted 100 SM backbone genes on average. Yet, the trends are similar as both studies report *Aspergillus floccosus* as the richest species in SM backbone genes within the *Terrei* section (Theobald et al., 2024). Five groups of SM backbone genes stood out due to their abundance in all genomes: NRPS-like with 3,888, rPKS (reducing Polyketide Synthase) with 3,175, NRPS with 2,943, nrPKS (non-reducing Polyketide Synthase) with 1,395, and DMATS with 1,203 backbone genes. In 145 species, and the three selected *Penicillium*, NRPS-like, rPKS, and NRPS are the three most abundant SM backbone genes. In contrast, ten species in sections *Flavipedes* and *Fumigati* have highly abundant DMATS, and other 22 Aspergilli have also many nrPKSs. Interestingly, we also observed variation in the number of SM backbone genes within sections. For example, within the *Nigri* section, two sub-groups could be defined based on their morphology, biseriates and uniseriates which differ in the attachment of phialides to the vesicle. Among them biseriates tend to have more SM backbone genes harbored in their genomes compared to uniseriates (Nielsen et al., 2009). Similarly, two sub-groups in the *Fumigati* section show a diverse pattern of SM backbones, the clade to which *A. fumigatus* belongs having a higher number of SM backbone genes. In summary, our findings underscore that the SM repertoires are diverse and particularly vary between sections.

Previous studies used BGC network approaches to assess BGC conservation within *Aspergillus* sections (Kjærbølling et al., 2020; Vesth et al., 2018), which overestimates the BGC diversity as variation in gene content at the loci of predicted BGCs will directly result in non-conserved BGCs. In contrast, we here utilized a phylogeny-aware method (Figure S1) to determine orthologous groups for each type of SM backbone gene. To obtain an estimate of gene conservation in Aspergilli, we used OrthoFinder to identify orthologous groups for all genes among all genomes. We included in total 1,546,732 orthologs as defined by OrthoFinder (Table S4) and 16,638 SM backbone genes to test their uniqueness among Aspergilli. From all OrthoFinder entries, we identified 34,144 unique genes (2.2%) among all species, for which the estimated growth rate β is 0.27 (Figure 2A), where β represents the rate of growth of unique genes as the number of sampled species increases (Heaps, 1978). For SM backbone genes, 2,426 out of 16,638 (14.6%) are unique, for which the growth rate β is 0.45 (Figure 2B). These β rates imply that sequencing more *Aspergillus* species is expected to expand the overall genomic diversity and specifically increase the diversity within the set of SM backbone genes. Interestingly, a recent study observed more diverse SM genes compared to pangenomes at the population level (Steenwyk et al., 2023), suggesting a large, yet untapped diversity of SMs in Aspergilli. However, in our study, saturation is reached for many SM groups, but not for NRPS and NRPS-like, which indicates that we might have found nearly all polyketide and terpene biosynthetic pathways, but many non-ribosomal peptides likely remain to be discovered. Lastly, we observed a similar trend for both unique orthologues and SM backbone genes with continuous slight increases within sections and small jumps between sections (Figures 2C and 2D), suggesting that sections are indeed characterized by a sudden increase in BGC diversity that is consistent with divergence.

**Figure 2.**
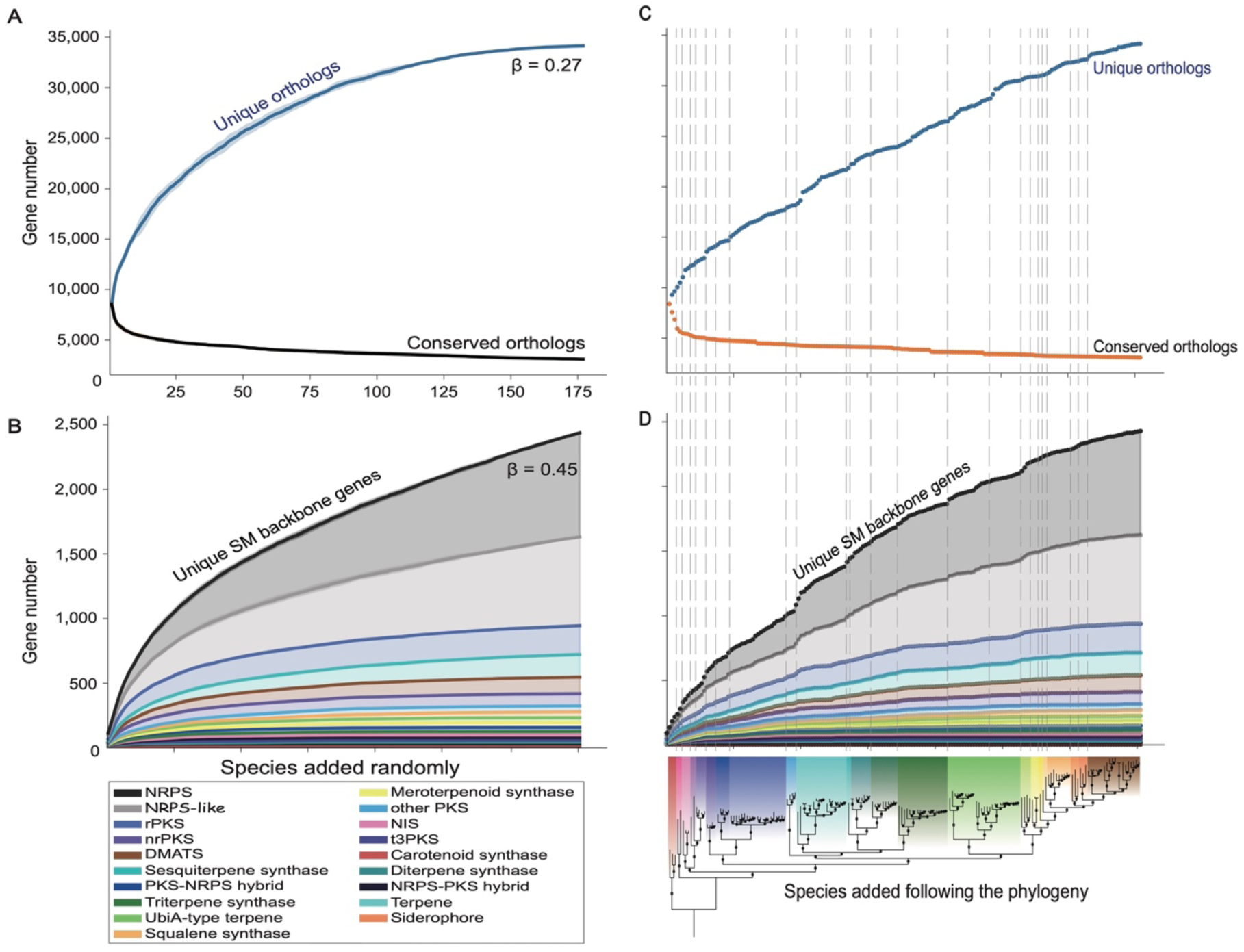
Evaluation of the repertoire of unique secondary metabolite (SM) backbone genes in Aspergilli. (A) The saturation curve plots the number of unique and conserved orthologous genes when species are added randomly. The calculations were performed ten times, and the standard deviations are shown as shadows around the curve. (B) Saturation curves plot the stacked number of unique SM backbone genes when species are added randomly. Different colors indicate distinct groups of SM backbone genes. The calculations were performed 10 times, and the standard deviations are shown as shadows around the curve. (C) Saturation curves of unique and conserved orthologous genes when species are added according to their taxonomy, starting with the outgroup, and the dash lines indicate distinct taxonomic sections. (D) Saturation curves of unique SM backbone genes with the species added according to their taxonomy and the dash lines indicate distinct taxonomic sections. A phylogenetic species tree is shown at the bottom for reference. PKS: polyketide synthase; NRPS: non-ribosomal peptide synthetase; DMATS: dimethylallyl tryptophan synthase; NIS: NRPS-independent siderophore.

We observed that SM backbone genes are diverse and that increased diversity is observed across different *Aspergillus* sections. However, the number of unique genes is lower than estimated in three recent studies focused solely on the *Nigri* (Vesth et al., 2018), *Flavi* (Kjærbølling et al., 2020) or *Terrei* (Theobald et al., 2024) sections that found around half of the SM BGCs were predicted to be species-specific. Here, we only focused on SM backbone genes and implemented a phylogeny-aware approach to evaluate SM backbone gene diversity across a diverse set of *Aspergillus* species, while these previous studies used complete SM gene cluster diversity and relied on a limited taxonomic sampling. These previous studies thus overestimated the BGC diversity within *Aspergillus* sections. In contrast, our results are more conservative and underestimate the true biochemical diversity that is provided by the diversity of additional tailoring genes within BGCs that can lead to the production of structurally related yet different secondary metabolites. However, our approach provides broader and more detailed estimations regarding the type of metabolite and BGCs that remain to be discovered in *Aspergillus* species.

### Genome-wide synteny in Aspergilli is higher within than between sections

The tight regulation of BGC expression and the role of chromatin conformation in this regulation suggested that the genomic localization of BGCs might be important (Palmer et al., 2010). Several studies have indicated an over-representation of SM BGCs in sub-telomeric regions (Guzmán-Chávez et al., 2018; Keller, 2019; Kjærbølling et al., 2020; Klejnstrup et al., 2012). These regions are known to be variable between strains and species, and consequently, they display low levels of synteny. The term ‘synteny’ here refers to the conservation of genomic regions that are maintained in a similar arrangement across different genomes through evolutionary time (Drillon et al., 2013) (Figure 3A). As most of the *Aspergillus* species do not have chromosome-level assemblies, we assessed BGC localization by examining synteny throughout the *Aspergillus* genus. We performed an all-versus-all pairwise comparison, and we uncovered, in general, that the amount of shared synteny between species (synteny percentage) correlates with the divergence and is consistent with the phylogenetic relationship between species (Figure 3B); *i.e*., two *Aspergillus* sections that are more syntenic indeed diverged more recently. Based on this observation, we propose that the *Nigri* section could be split in two, which is also coherent with the phylogeny. Similarly, the *Circumdati*, *Flavi*, and *Nidulantes* sections show distinct groups of higher synteny that might indicate ongoing divergence within these sections. Section *Nidulantes* exhibits lower synteny, which seems coherent with the branch lengths in this clade compared to other clades in which the divergence between species appears more recent than in the *Nidulantes* section. Finally, early-diverging sections (*i.e., Polypaecilum*, *Cremei*, *Restricti*, *Aspergillus*, *Cervini*, *Clavati*, and *Fumigati*) show high synteny within sections compared with other sections (Figure 3C). Overall, the observed synteny between sections is low, ranging from only 34% to 54%, with a median of 46%. The same analysis performed with only highly contiguous assemblies yielded the same results (Figure S2B) (Liu et al., 2018).

**Figure 3.**
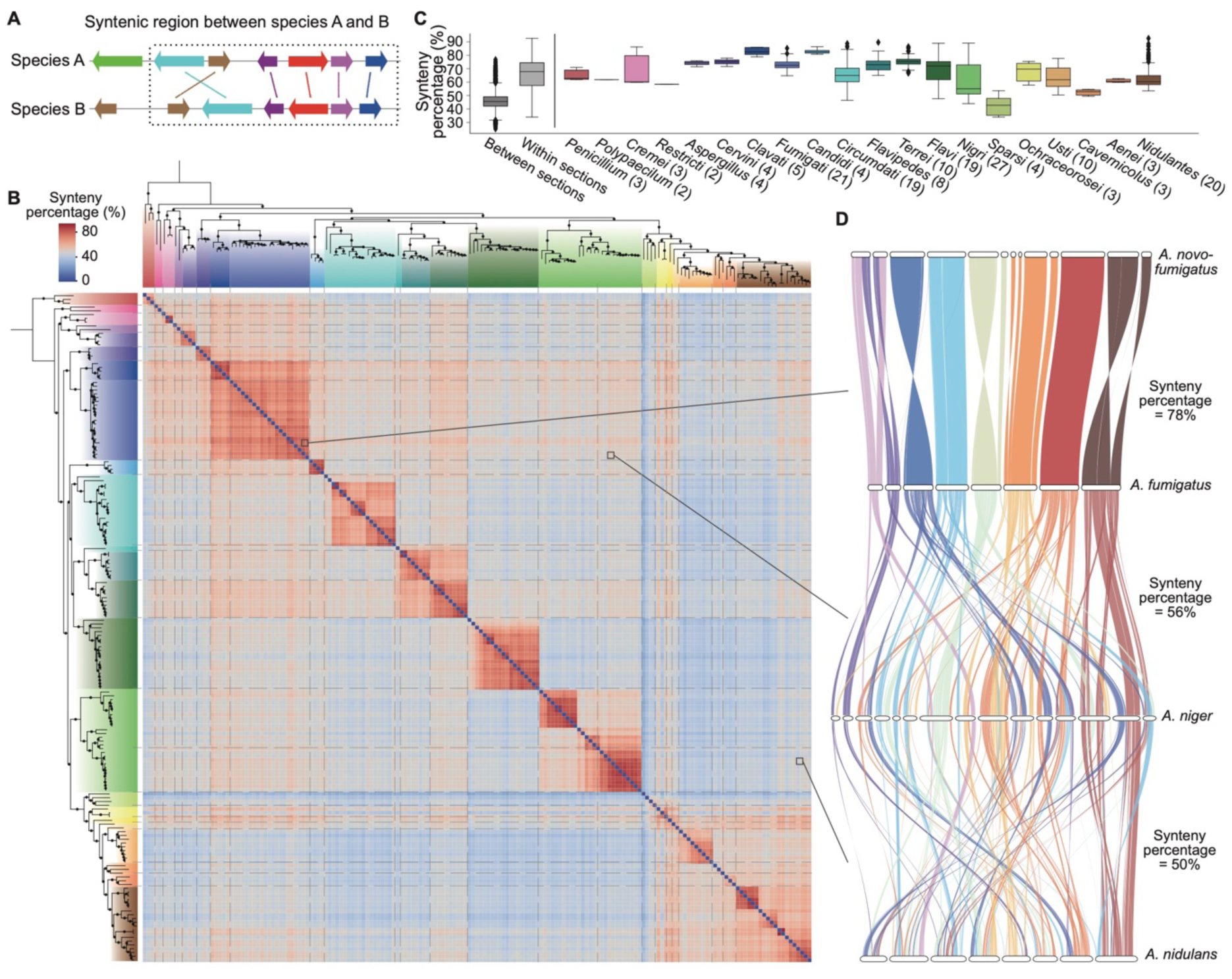
Conservation of synteny within and between *Aspergillus* sections. (A) A graphical representation of a hypothetical syntenic region between two species is demarcated by a dashed box. Arrows of the same color, linked by a line, represent homologous genes from two different species, with the arrow direction indicating the gene’s orientation. (B) The heatmap shows pairwise synteny expressed as the synteny percentage based on all-versus-all comparisons, arranged based on the phylogenetic tree (Figure 1). The species tree, displayed to the left and top of the heatmap, uses colors to differentiate between sections (see legend for Figure 1 for the color coding). (C) Boxplots show the quantification of synteny by the ‘synteny percentage’ between sections, within sections, and in each section of the *Aspergillus* genus. Numbers in the parentheses indicate the number of species in the sections, and only sections with more than one species are shown. (D) Genome alignment between two species from the same section (top) and different sections (middle and bottom) highlights the degradation of synteny when taxonomic distance increases.

In Ascomycota, synteny varies across distantly related species and is categorized into micro-, meso-, and macro-synteny (Hane et al., 2011). Micro-synteny refers to the conservation of a small number of successive genes, typically ranging from two to ten. Meso-synteny represents larger, yet fragmented, synteny blocks compared to micro-synteny, but shorter than 20 thousand base pairs (kbp). Macro-synteny, on the other hand, is characterized by large and continuous blocks extending over 20 kbp (Hane et al., 2011). Our findings align with the expectations, revealing longer synteny blocks within sections than between them (Figure S2). For instance, species in the *Fumigati* section show extensive regions of shared synteny across chromosomes (Figure 3D). In contrast, species from different sections exhibit markedly reduced synteny, both in percentage and block length (Figures 3D and S2). A pattern of ‘degraded macrosynteny’ was previously identified in several filamentous Ascomycota fungi, including some Aspergilli (Hane et al., 2011). Our data suggest that within *Aspergillus* sections, synteny blocks with an approximate length of 150 kbp are indicative of macrosynteny. In contrast, between different sections, synteny blocks averaging 70 kbp to 150 kbp represent degraded macrosynteny, likely reflecting the divergence time among these sections.

### SM backbone genes tend to reside in low-synteny and sub-telomeric regions

To investigate the genomic localizations of SM genes in Aspergilli, we analyzed four species with chromosomal-level assemblies: *A. nidulans*, *A. fumigatus*, *A. niger*, and *A. oryzae* (Figures 4 and S3-S5). In each of the four species, the gene density shows that predicted genes are evenly distributed throughout the genome, except for putative centromere and telomere regions (as defined by the terminal 10% regions of chromosomes), which are known to contain fewer genes but harbor repetitive sequences, and exhibit unique structural and functional characteristics distinct from the rest of the genome (Xu et al., 2022). BUSCO genes, which are conserved across the fungal kingdom and crucial for organismal function, are significantly enriched in regions with high synteny, both within individual sections (91% of BUSCO genes; Chi^2^ *p*-value = 3.15^e-117^) and across the *Aspergillus* genus (83%; Chi^2^ *p*-value = 0) compared with all genes (59% and 42%, respectively) (Figure 5B and 5C). In contrast, SM backbone genes are significantly enriched in low-synteny regions in all four species at the *Aspergillus* genus level (81% of backbone genes; Chi^2^ *p*-value = 6.12^e-58^) (Figures 5B and 5C, Table S6). For the section-level synteny scores, though only 24% of SM backbone genes are found in low-synteny regions, the trend remains significant compared with all genes (14%; Chi^2^ *p*-value = 7.77^e-28^) and clearly differs from BUSCO genes (2%) (Figures 5B and 5C, Tables S6). In regard to chromosomal location, only 21% of all genes are located in sub-telomeric regions, and BUSCO genes are significantly located outside of these regions (7% in sub-telomeric regions; Chi^2^ *p*-value = 4.34^e-72^) (Figures 5D and 5E). In contrast, SM backbone genes show a trend to be located in the sub-telomeric regions (36%; Chi^2^ *p*-value = 1.28^e-12^) (Figures 5D and 5E), aligning with previous reports in *A. nidulans* (Klejnstrup et al., 2012), *A. fumigatus* (Keller, 2019; Perrin et al., 2007), *A. oryzae* (Kjærbølling et al., 2020), as well as in other fungal species such as *F. fujikuroi* (Studt et al., 2017) and *Penicillium chrysogenum* (Guzmán-Chávez et al., 2018). However, the enrichment of SM backbone genes in sub-telomeric regions is less significant than in low-synteny regions, suggesting that association with low-synteny regions is an attribute of SM BGCs. The localization of SM backbone genes in low-synteny and sub-telomeric regions of *Aspergillus* genomes may facilitate rapid genetic rearrangement of these BGCs, thereby enabling rapid adaptation to diverse ecological niches. Indeed, such a localization is coherent with the observed isolate- or population-specific presence-absence pattern reported in *A. nidulans* (Drott et al., 2020), *A. fumigatus* (Lind et al., 2017) and *A. flavus* (Drott et al., 2021). Most polymorphisms detected in *A. fumigatus* population were suggested to involve transposable elements and/or recombination (Lind et al., 2017), which are mechanisms likely to break synteny and could favor this bias in BGC genomic localization. Noteworthy, SM backbone genes with known metabolite structures, as listed in the MIBiG database (Table S5), tend to occur more frequently in sub-telomeric areas. For instance, four out of six SM backbone genes in *A. nidulans* (Figure 4), 18 out of 27 in *A. fumigatus* (Figure S3), and seven out of eleven in *A. niger* (Figure S4) are found in sub-telomeric regions. *A. oryzae* represents a unique case, with only two out of 18 SM backbone genes with known structures located in sub-telomeric regions (Figure S5). The observation that most characterized BGCs are localized in sub-telomeric regions suggests that they are more frequently expressed than those in other regions.

**Figure 4.**
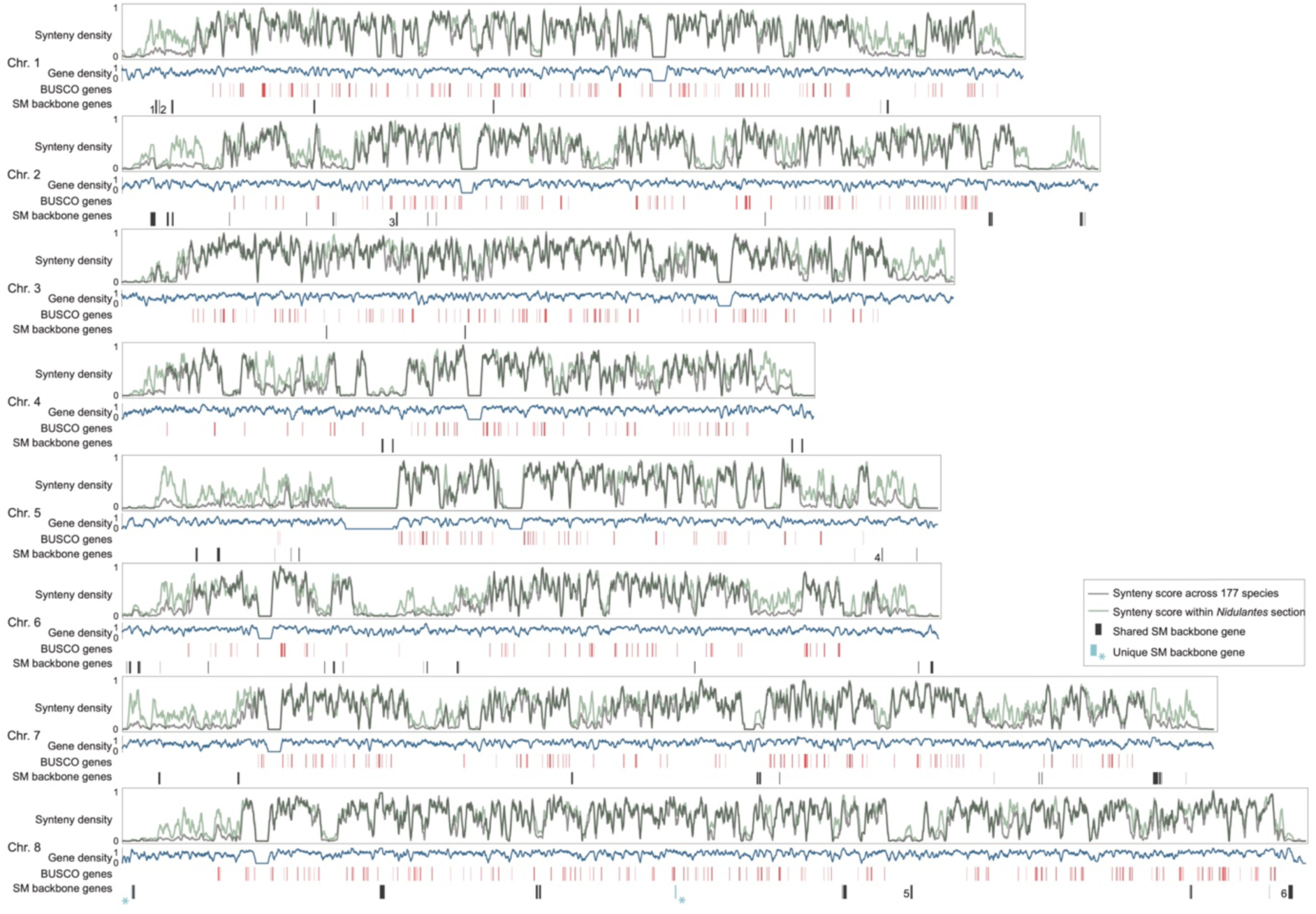
Chromosomal localization of secondary metabolite (SM) backbone genes in *Aspergillus nidulans*. This figure illustrates the genomic localization of BUSCO genes and SM backbone genes in *Aspergillus nidulans*. Lines in black and green represent synteny density across 177 species and within the *Nidulantes* section, respectively. The gene density is shown along the chromosomes. Red bars indicate the chromosomal locations of BUSCO genes, with bar width proportional to gene length. Black and blue bars mark the positions of shared and unique SM backbone genes, respectively, with their widths corresponding to gene length. Numbers adjacent to the black bars denote SM backbone genes associated with known metabolites as listed on the MIBiG website (Table S5).

**Figure 5.**
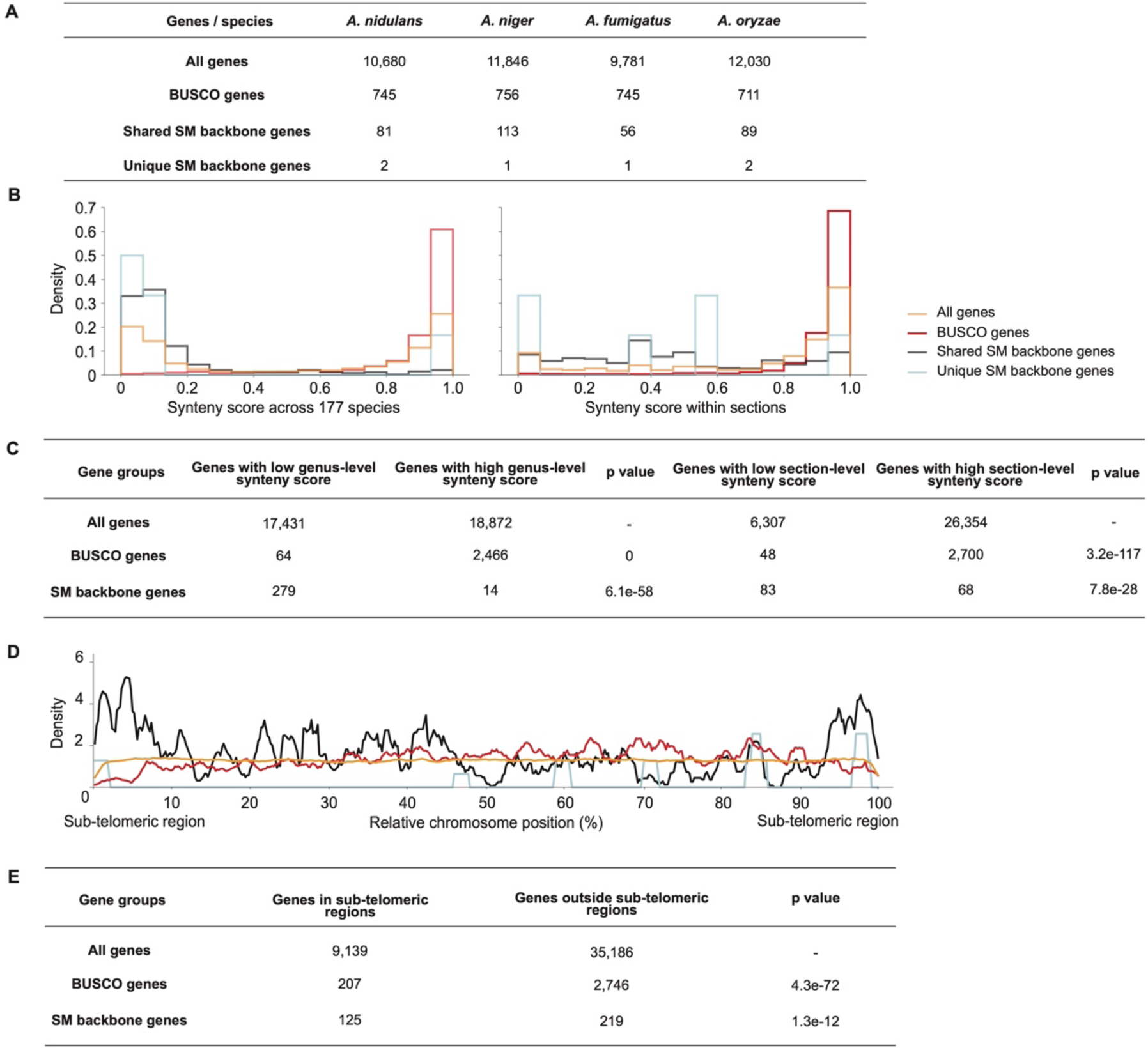
Analysis of secondary metabolite (SM) backbone gene localization in low-synteny and sub-telomeric regions in four *Aspergillus* species. (A) Summary table of the different groups of genes considered in four *Aspergillus* species. (B) Distribution of different groups of genes in relation to synteny score analyzed across 177 species and within individual sections. (C) Summary table of the number of SM backbone genes, BUSCO genes, and all genes with high-synteny score (larger than 80%) and low-synteny score (lower than 20%) at the genus-level and section-level. (D) Distribution of different groups of genes relative to chromosomal localization, considering the terminal 10% of chromosomal regions as sub-telomeric. (E) Summary table of the number of SM backbone genes, BUSCO genes, and all genes within and outside of sub-telomeric regions. Chi-square statistic tests (Virtanen et al., 2020) were performed on SM backbone genes and BUSCO genes, determining whether their distributions differ significantly from the distribution of all genes in terms of genus-level synteny, section-level synteny, and sub-telomeric localization.

### Exploring the impact of genomic localization and histone PTMs on SM gene expression variability in Aspergilli

It has been proposed that genomic localization of SM BGCs is important for the regulation of their expression through histone modifications as several histone PTMs were shown to influence SM production in Aspergilli (Cánovas et al., 2014; Lee et al., 2009; Li et al., 2019). Thus, we sought to explore the relationship between gene expression, histone PTMs, and genomic and syntenic localization of SM backbone genes compared with BUSCO genes. To this end, we utilized publicly available RNA-seq data from *A. nidulans*, *A. fumigatus*, *A. niger*, and *A. oryzae*, and ChIP-seq (Chromatin ImmunoPrecipitation Sequencing) data from the first two Aspergilli (Tables S7 and S8). We used the coefficient of variance (CV) to assess the relative variability of gene expression and histone modification of different gene groups across diverse experimental conditions. For BUSCO genes, we observed stable gene expression in all species regardless of the synteny score (Figures 6A and S6), which is accompanied by limited variation in all histone PTMs included in this study (H3K4me3, H3K9me3, H3K36me3, and H3Kac) (Figures 6B-E, S6, S7). Compared to BUSCO genes, SM backbone genes exhibit greater variability in gene expression (Figures 6A and S6). For histone PTMs, H3K36me3 in *A. nidulans* and H3K4me3 in both *A. nidulans* and *A. fumigatus* show higher variability at SM backbone genes compared with other gene groups (Figures 6B-C, S7 and S8), suggesting a role for these histone modifications in regulating the expression of SM genes. Especially, SM backbone genes with known structure and variable expression in *A. fumigatus* also exhibit high CVs in H3K4me3, with four of them ranking among the top five most variable SM genes (Table S5). However, the observed gene expression variation of SM genes is not explained by H3K9me3 (in *A. nidulans* and *A. fumigatus*) nor H3ac (in *A. nidulans*) as SM genes do not differ from BUSCO genes for these PTMs (Figures 6D-E, 6H-I, and S8). The finding that gene expression seems independent from H3ac contrasts with the results obtained with *A. nidulans* histone deacetylase mutants in which the expression of many BGCs is affected (Lee et al., 2009). A possible explanation is that the inability to deacetylate histone tails prevents silencing through histone methylation. It also suggests that specific histone methylations are a stronger determinant of BGC expression than their acetylation. When comparing to other accessory genes, as defined by genes located in regions with a synteny score lower than 0.2, SM genes do not seem to differ in regard to RNA CVs and ChIP-seq CVs, indicating that the genomic localization is particularly important for the regulation of the expression.

**Figure 6.**
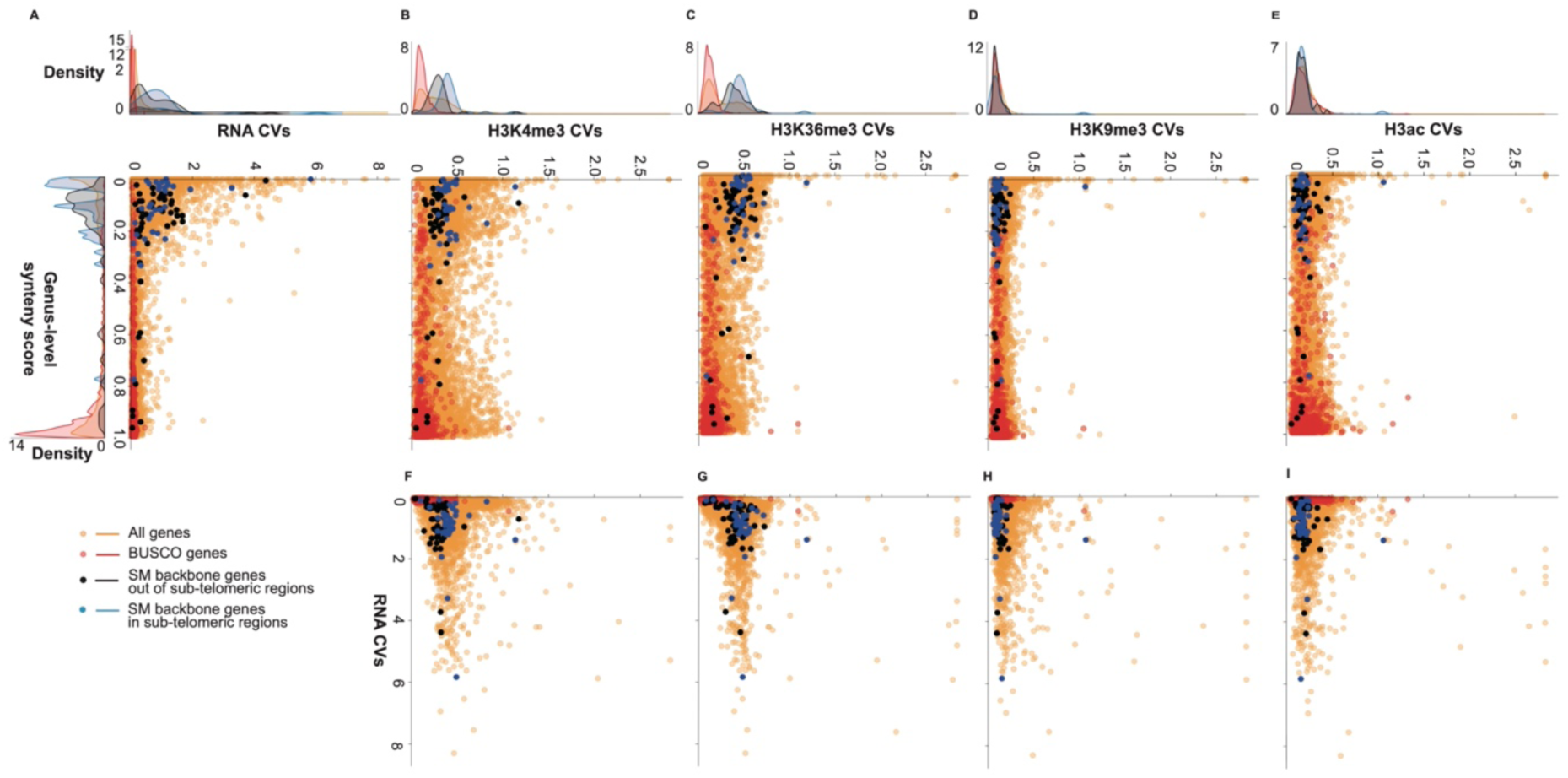
Correlation between genomic localization of secondary metabolite (SM) backbone genes and variation in gene expression and histone modifications in *Aspergillus nidulans*. We employed the coefficient of variance (CV) to assess the relative variability in gene expression (RNA) and histone post-translational modifications of all genes, BUSCO genes, and SM backbone genes in *Aspergillus nidulans*. (A-E) Correlation between CVs and genus-level synteny. (F-I) Correlation between histone modification CVs and gene expression CV. Me3: trimethylation; ac: acetylation.

When taking the chromosome localization into account, SM backbone genes within the sub-telomeric regions show slightly higher variation in gene expression in *A. nidulans*, *A. fumigatus*, and *A. niger*, but not in *A. oryzae* (Figures 6A and S6). The difference in *A. oryzae* could be due to the fact that it was domesticated for industrial purposes such as producing sake, soy sauce, and miso, and it is generally recognized for its simplified metabolic profile and inability to produce aflatoxins (Gibbons et al., 2012). A similar influence of domestication on the downregulation of metabolite production was also observed in wild *Penicillium* molds on cheese (Bodinaku et al., 2019). However, the sub-telomeric localization does not fully explain a higher variation in both gene expression and histone PTMs because the distribution of these variations for SM backbone genes outside sub-telomeric regions is bimodal and overlap with sub-telomeric backbone genes (Figures 6A, 6C, S6, and S7). Lastly, SM backbone genes with the known metabolite structures listed on the MIBiG website show various expression patterns (Table S5). In *A. fumigatus*, known SM genes show high variability as their average CV (1.20) is higher than the average CV for both all SM backbone genes (1.09) and SM backbone genes outside sub-telomeric regions (0.94) (Table S5). Two SM backbone genes responsible for sartorypyrone A, located in a low-synteny region at the genus level but not in a sub-telomeric region (genes 25-27 in Figure S3), are in the top five most variably expressed SM backbone genes. By contrast, in other Aspergilli, either no (in *A. nidulans* and *A. niger*) or only one (in *A. oryzae*) SM backbone gene with known structure is in the top five most variable SM backbone genes (Table S5). Based on our results, SM backbone genes that are still uncharacterized but exhibit higher variation in expression deserve further attention as they are likely to produce SMs under laboratory conditions. It must be noted that our conclusions rely on a very limited set of available RNA-seq and ChIP-seq data. Gene expression and histone PTMs are highly context-dependent, and the calculated CVs could be biased. We recommend that future studies aim to incorporate more histone PTMs and expression data from diverse conditions to mitigate these limitations and enhance the robustness of the conclusions drawn.

### Conclusions

Altogether, our results highlighted that SM backbone genes are quite abundant and diverse in the fungal genus *Aspergillus*, especially between different sections. Moreover, based on the four *Aspergillus* species with chromosomal-level assemblies and publicly available RNA-seq and ChIP-seq data, we showed that SM backbone genes exhibit a strong bias for low-synteny regions, and the association between their localization, variation in expression, and histone PTMs is restricted to H3K4me3 and H3K36me3 only. It is important to note that our study is limited to the few histone PTMs that are publicly available and other PTMs might play a more important role and need to be investigated. In particular, H3K79me is a relevant candidate as this modification was reported to localize at transition regions between heterochromatin and euchromatin, and we recently reported for this modification differences in abundances between different Aspergilli (Zhang et al., 2023). While our understanding of how genomic localization and histone PTMs regulate SM production remains partial, our study points at new research directions on understudied histone modifications that might prove key to unlock the full potential of SM production in Aspergilli.

## Materials and methods

### Acquisition of genomes and predicted proteomes for 177 fungal species

The genomes and predicted proteomes of 177 fungal species were retrieved from the Joint Genome Institute MycoCosm (https://mycocosm.jgi.doe.gov/mycocosm/home) repository (Grigoriev et al., 2014) in January 2022. These species comprise 174 Aspergilli as well as three *Penicillium* species (*Penicillium brevicompactum*, *Penicillium griseofulvum*, and *Penicillium chrysogenum*) that were used as an outgroup in our analyses (Table S1).

### Species tree reconstruction

To reconstruct the phylogenetic relationship between the here analyzed *Aspergillus* species, we used the fungal database (Fungi Odb10, 758 orthologs) in BUSCO (Benchmarking Universal Single-Copy Ortholog assessment tool) v4.0.1 (Simão et al., 2015; Zdobnov et al., 2021) to retrieve fungal single-copy genes (Table S2) that are present in the 177 genomes. For homologs of each single-copy BUSCO gene, we first generated multi-sequence alignment using MAFFT v7.271 (parameter: --auto) (Katoh & Standley, 2013), then removed positions in the alignments with gaps in more than 90% of the sequences using ‘-gt 0.1’ in TrimAl v1.2 (Capella-Gutiérrez et al., 2009) for better tree construction (Steenwyk et al., 2020). We concatenated the 758 alignments into one ‘super-alignment’ and employed IQ-TREE v1.6.10 (Chernomor et al., 2016; Nguyen et al., 2015) to construct the maximum-likelihood phylogeny with partitioned model selection to allow selecting a substitution model (Table S2) for each BUSCO gene alignment, and used both ultrafast bootstrap approximations and SH-aLRT (Shimodaira–Hasegawa approximate Likelihood Ratio Test) with 1,000 pseudo-replicates to calculate branch supports (Hoang et al., 2017).

### Identification and conservation analysis of putative SM backbone genes and orthologous groups

We used antiSMASH (antibiotics and Secondary Metabolite Analysis SHell) v6.1.0 (Blin et al., 2021) (parameters: –minimal) to identify all regions with predicted BGCs in the 177 genome assemblies, and refined the classification of SM backbone genes with our in-house tool BGCtoolkit (https://github.com/WesterdijkInstitute/BGClib) based on conserved domain organization (Figure 1 and Table S3). Then, we extracted the protein sequence of the predicted backbone enzymes or of the adenylation domains for NRPS, NRPS-like, and PKS-NRPS hybrid enzymes using BGCtoolkit. For each type of backbone enzyme, protein sequences were aligned and trimmed using MAFFT and TrimAl, respectively, as mentioned above. We constructed phylogenetic trees using FastTree v2.1.11 with default parameters (Price et al., 2009). We subsequently used Newick Utilities v1.6.0 (Junier & Zdobnov, 2010) to identify orthologs *via* three steps: i) re-root the tree based on the mid-point (nw_reroot); ii) manually select the clades having more than 88 leaves and with trustworthy bootstrap value (>= 0.99) and mark them as orthologs; iii) for the remaining leaves, automatically split the clades with fewer than 88 leaves and with trustworthy bootstrap value (>= 0.99) (nw_ed -n -r). In the construction of the three adenylation domain trees, an extra step was undertaken to consolidate multiple domains originating from the same gene. This involved examining all leaves in the domain tree to identify the genes associated with these domains. Clades were collapsed and assigned a new clade name when the domains were found to originate from the same gene. For any remaining domains, the original clade names were retained. We built a saturation curve with the matplotlib.pyplot visualization library (Hunter, 2007) and used the function from the SciPy library (Virtanen et al., 2020) to implement a curve-fitting process, which determines the rate of growth (β) of unique gene numbers under Heap’s Law (Heaps, 1978).

We used OrthoFinder v2.3.8 (Emms & Kelly, 2019) to identify orthologous groups across 177 predicted proteomes using default settings (Table S4). A saturation curve based on these orthologous groups was built as described above.

### Genome synteny analysis

We employed MCScanX (Multiple Collinearity Scan Toolkit X version) (Wang et al., 2012) to conduct an all-versus-all pairwise comparison among all genomes to identify syntenic blocks (match size: -s 3 and max gaps: -m 15). We defined the ‘synteny percentage’ between a pair of species as the ratio of syntenic genes detected between two compared species to the total genes. We defined the ‘between sections’ synteny percentage by considering all pairs of species that are associated with different taxonomic sections, and the ‘within sections’ synteny percentage by considering all pairs from the same taxonomic section. For the synteny percentage of each taxonomic section, we considered all comparisons for species within that specific section. For visual representation of genome alignment between two species, we used the R package GENESPACE v1.2.3 (Lovell et al., 2022).

To calculate the average length of syntenic regions, we took all syntenic blocks identified by the MCScanX in all-versus-all pairwise comparisons and excluded the contigs with less than 50 genes. Subsequently, we employed the hypergeometric distribution (Rice, 2003) utilizing the ‘hypergeom.cdf(x, M, n, N)’ function from the SciPy library (Virtanen et al., 2020) to evaluate the likelihood of successfully detecting a syntenic region between two given species. In this function, ‘M’ denotes the total number of homologous gene pairs identified when comparing the complete genomes of species A and B, ‘n’ represents the count of genes on scaffold B1 with homologs in species B, and ‘N’ is the number of genes on scaffold A1 with homologs in species A. This cumulative distribution function aggregates the probabilities of identifying up to ‘x’ successful syntenic matches, thereby yielding the cumulative probability of detecting ‘x’ or fewer successes. The analysis considered (1) the total number of homologs between the two species, (2) the number of homologs on each contig relative to the other species, and (3) the number of homologs between the contigs themselves. Setting a p-value threshold at 1e-5, we then computed the average length of syntenic blocks for each pair of species.

We defined the synteny score at the genus level as the synteny score for a specific gene by dividing the number of species containing that gene by the total number of species (177 species). At the section level, the synteny score for a gene is determined by the ratio of the number of species within the section harboring that gene to the total species count in that section. We used the function ‘chi2_contingency’ from the SciPy library (Virtanen et al., 2020) to perform chi-square statistic tests on SM backbone genes and BUSCO genes, determining whether their distributions differ significantly from the distribution of all genes in terms of genus-level synteny, section-level synteny, and sub-telomeric localization (Table S6).

### RNA-seq sample preparation

To perform RNA sequencing, we obtained the fine powder of young mycelium of *A. fumigatus* (strain: Af293, CBS 126847; grown on malt extract agar), *A. nidulans* (strain: Wtpaba (genotype: pabaA1); grown on yeast agar glucose), and *A. niger* (strain: NRRL3, CBS 120.49; grown on malt extract agar) as described previously (Zhang et al., 2023). Subsequently, we extracted high-quality total RNA from each species in four replicates, following our in-house RNA extraction protocol. Briefly, we placed around 100 mg of fine powder in a 1.5 mL tube, added 1 mL TRIzol reagent (Fisher, 15596026), and incubated for 5 mins at 25°C. Following that, we added 0.2 mL chloroform-d (Sigma, 416754), mixed gently by hand 15 times, and incubated for 5 mins at room temperature. After centrifuging the samples at 10,000 RPM for 15 mins, we transferred the aqueous supernatant to a new tube. We then added 0.5 volume of 100% ethanol (BOOM, 200-578-6) and proceeded from the 5^th^ step of the NucleoSpin RNA Kit protocol (Bioke, 740955). Finally, we eliminated DNA contamination using DNase I (Sigma, AMPD1) and sent samples to the Beijing Genome Institute (BGI, Hong Kong, China) for library construction and DNBseq (DNA NanoBall sequencing).

### Acquisition and analysis of public RNA-seq and ChIP-seq datasets

To retrieve publicly available RNA-seq data (Table S7) from *A. niger*, *A. fumigatus*, *A. nidulans*, and *A. oryzae*, we utilized prefetch v2.10.0 and fasterq-dump v2.10.0 from SRA Toolkit (https://github.com/ncbi/sra-tools) to extract FASTQ files from the SRA database (https://www.ncbi.nlm.nih.gov/sra). For all reads from both our BGI sequencing projects and the public database, we used the pseudoalignment tool kallisto v0.46.0 (Bray et al., 2016) to build the index and quantify raw read counts per gene. To remove the potential batch effect, we used ComBat-seq (Zhang et al., 2020) in R v3.6.1 (Team, 2010) and obtained batch-effect corrected read counts. Following this, we calculated the TPM (Transcripts Per Million) and used log-transformed TPMs to calculate the coefficient of variation (CV) to assess the variability of gene expression levels across different samples. We used seaborn (Waskom, 2021) and Matplotlib (Hunter, 2007) to plot the scatter plot and density plots.

We utilized ChIP-seq data from two studies (Colabardini et al., 2022; Gacek-Matthews et al., 2016), which included *A. nidulans* (H3ac, H3K4me3, H3K9me3, and H3K36me3) and *A. fumigatus* (H3K4me3 and H3K9me3) (Table S8). The analyses (downloading and processing) of this data were the same as the RNA-seq analysis mentioned above, yet we did not perform the batch effect removal step because the two studies were conducted in different species.

---

**Supplementary Figure 1.**
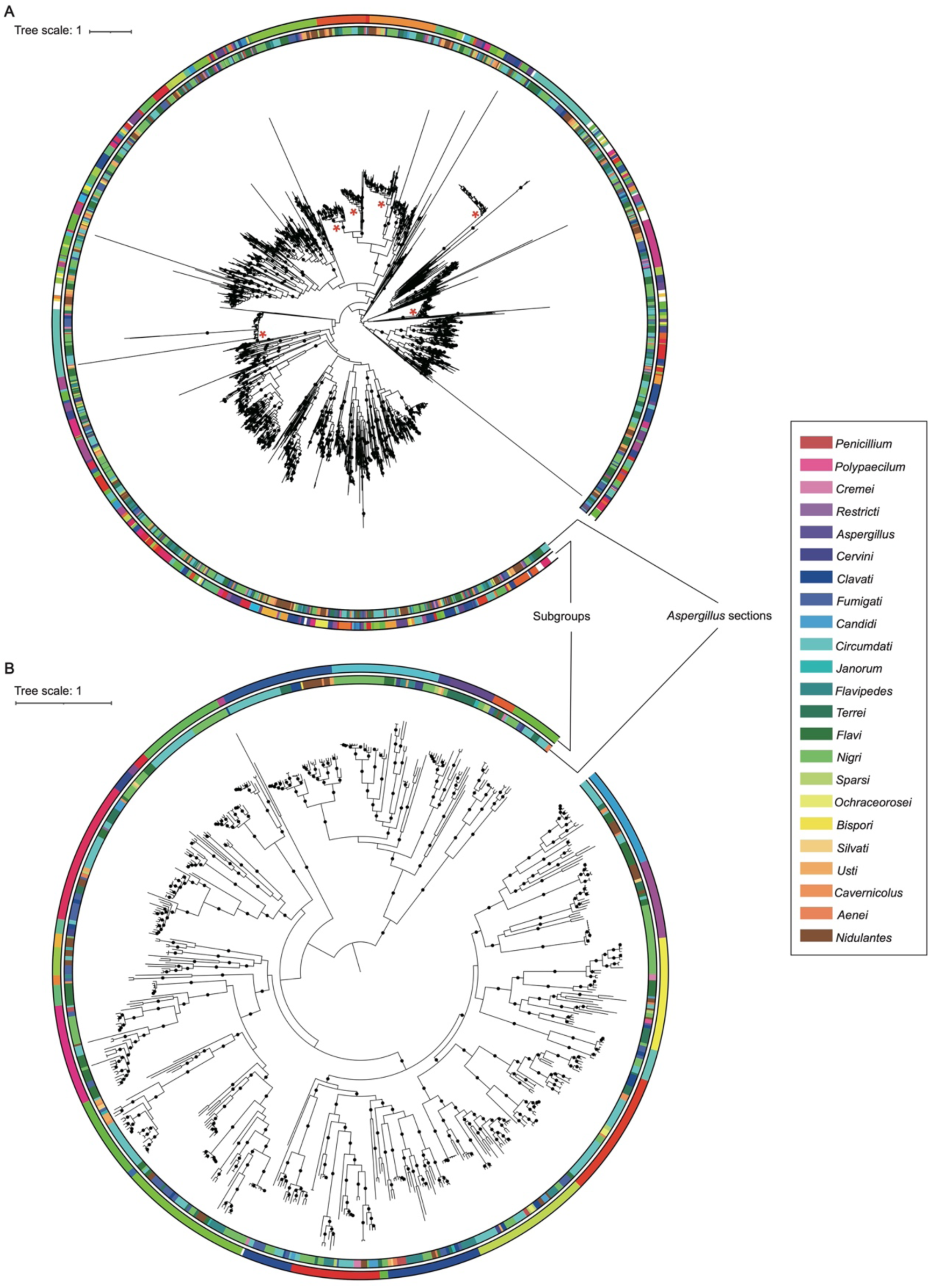
The phylogeny-aware method to detect secondary metabolite (SM) backbone gene diversity. We used FastTree (Price et al., 2009) to construct the phylogeny of each type of SM backbone genes and annotated the trees with iTOL. The inner colored ring indicates *Aspergillus* sections and the outer colored ring showed the splitting results by Newick Utilities (Junier & Zdobnov, 2010). (A) Manual selection is implemented first to pick up the clades larger than 88 leaves (red “*” in the tree). For remaining leaves and trees that do not need manual trimming (B), automatic splitting was performed with a trustworthy bootstrap value of >= 0.99.

**Supplementary Figure 2.**
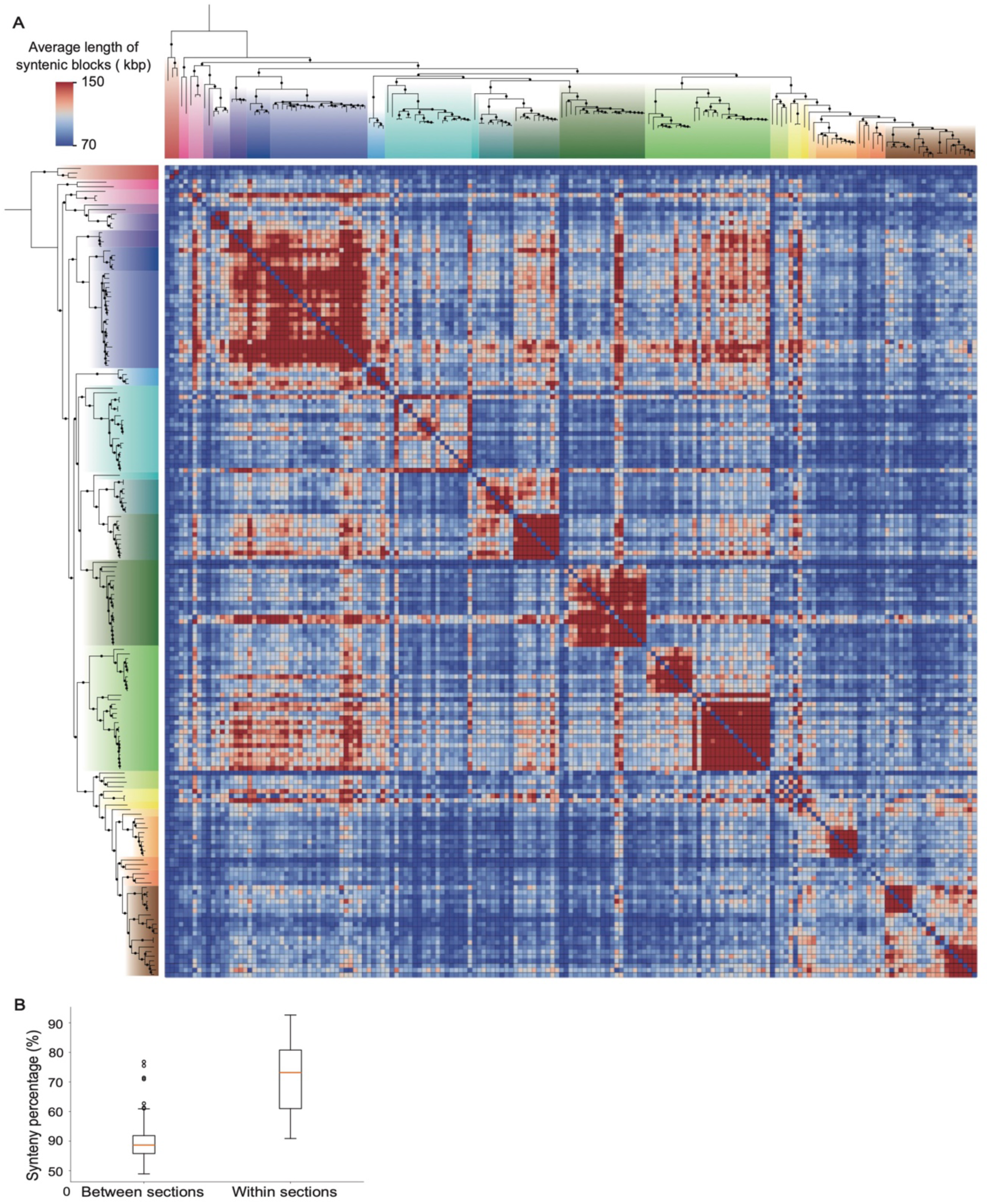
Length of syntenic blocks across *Aspergillus* sections. (A) The heatmap displays the all-versus-all pairwise comparisons based on the average length of syntenic blocks across all species; species were arranged according to the phylogenetic tree. The accompanying species tree, displayed to the left and top, uses colors to differentiate between sections. (B) We used the “highly contiguous” genome assemblies defined by (Liu et al., 2018) to calculate the synteny percentage between or within sections and obtained similar results as Figure 3C.

**Supplementary Figure 3.**
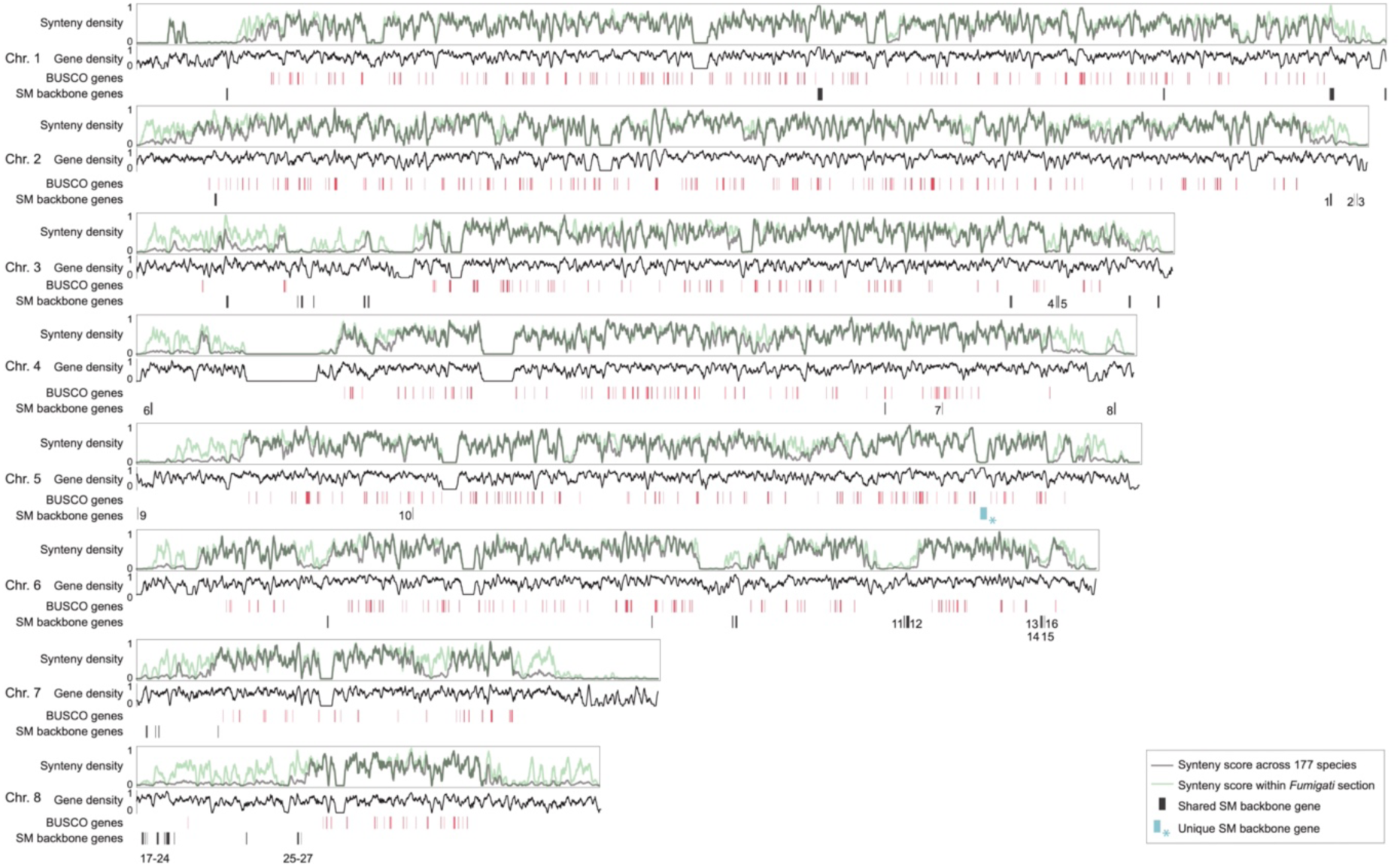
Chromosomal localization of secondary metabolite (SM) backbone genes in *Aspergillus fumigatus*. This figure illustrates the genomic localization of BUSCO genes and SM backbone genes in *Aspergillus fumigatus*. Lines in black and green represent synteny density across 177 species and within the *Fumigati* section, respectively. The gene density is shown along the chromosomes. Red bars indicate the chromosomal locations of BUSCO genes, with bar width proportional to gene length. Black and blue bars mark the positions of shared and unique SM backbone genes, respectively, with their widths corresponding to gene length. Numbers adjacent to the black bars denote SM backbone genes associated with known metabolites as listed on the MIBiG database (Table S5).

**Supplementary Figure 4.**
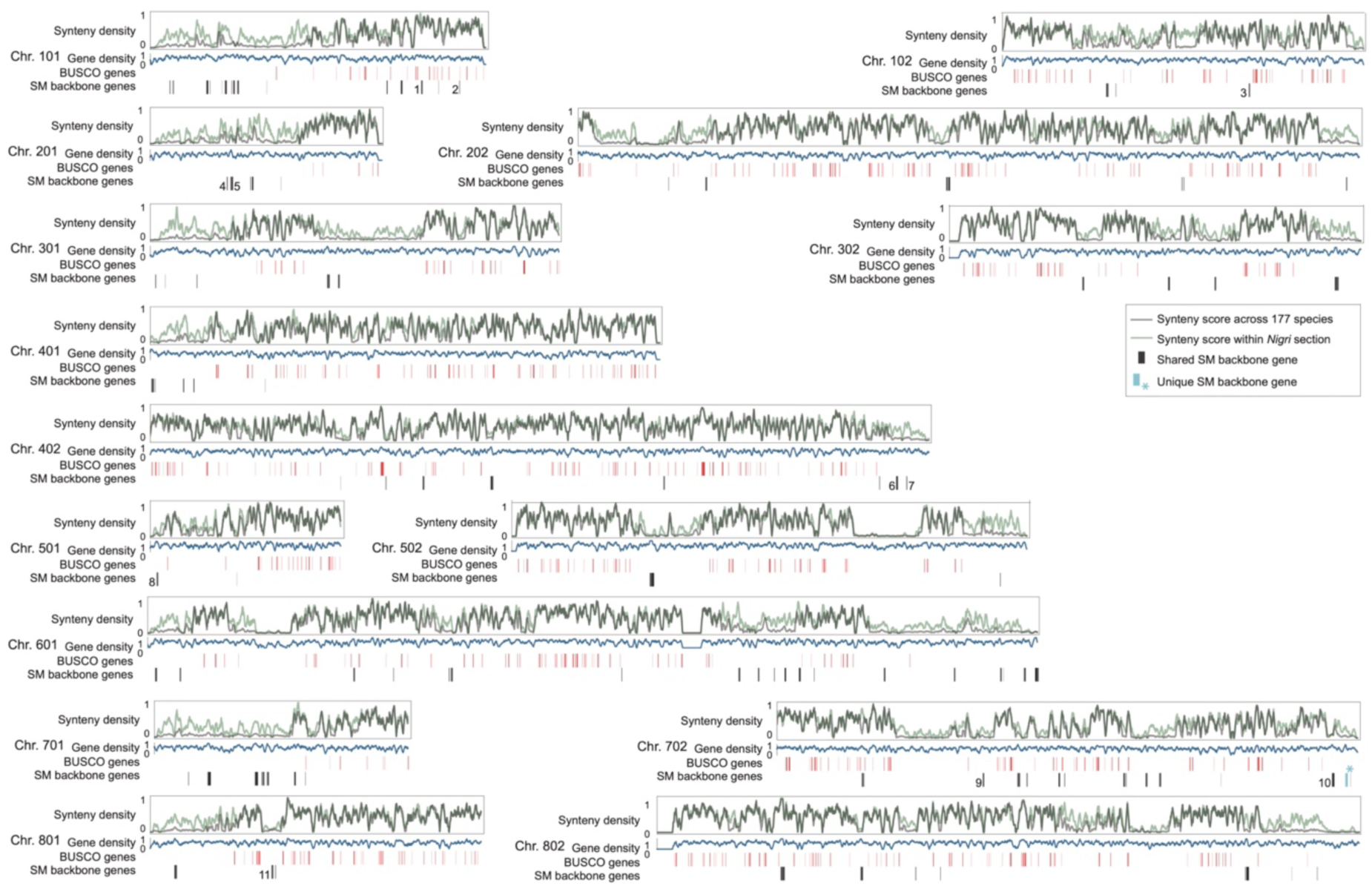
Chromosomal localization of secondary metabolite (SM) backbone genes in *Aspergillus niger*. This figure illustrates the genomic localization of BUSCO genes and SM backbone genes in *Aspergillus niger*. Lines in black and green represent synteny density across 177 species and within the *Nigri* section, respectively. The gene density is shown along the chromosomes. Red bars indicate the chromosomal locations of BUSCO genes, with bar width proportional to gene length. Black and blue bars mark the positions of shared and unique SM backbone genes, respectively, with their widths corresponding to gene length. Numbers adjacent to the black bars denote SM backbone genes associated with known metabolites as listed on the MIBiG website (Table S5).

**Supplementary Figure 5.**
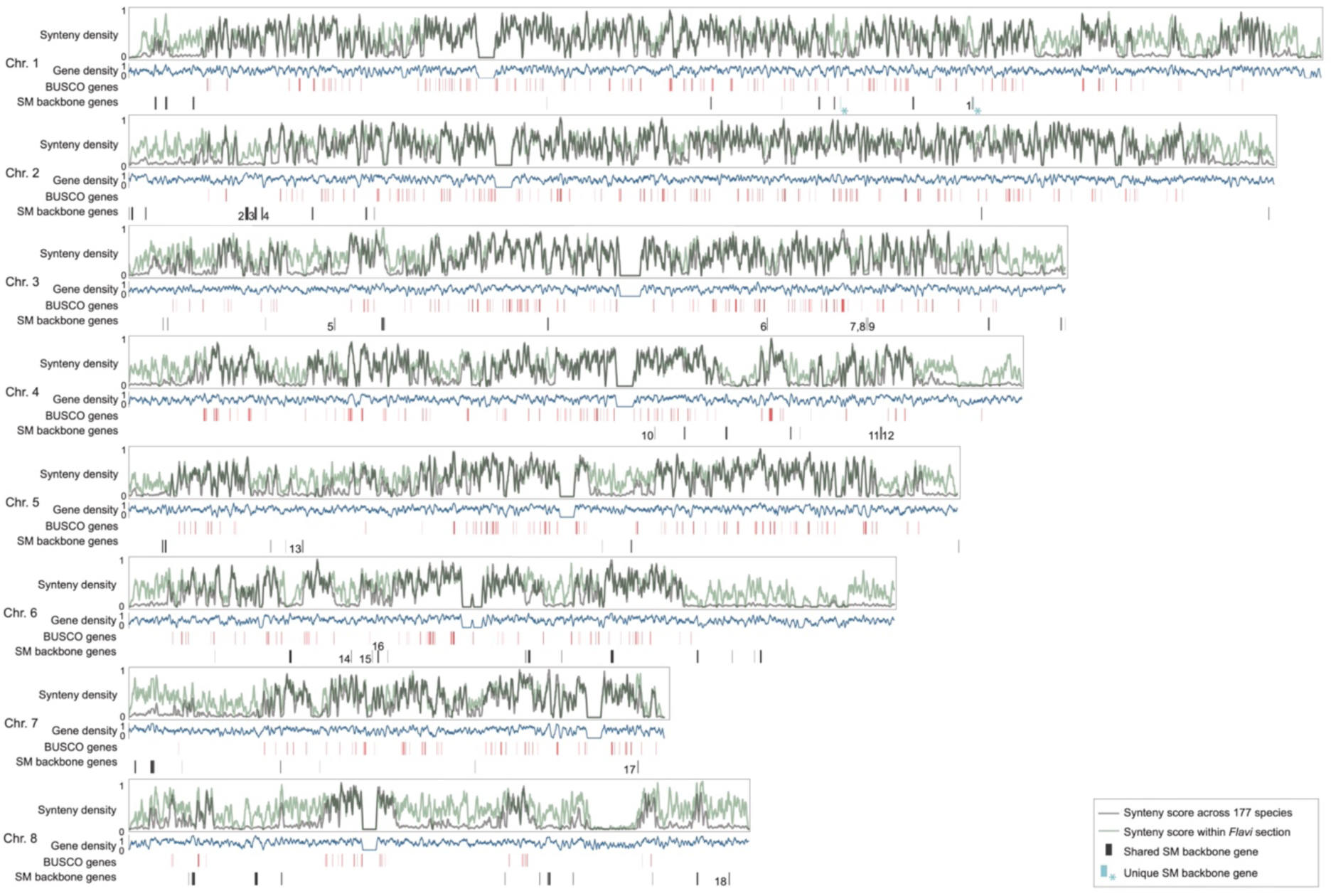
Chromosomal localization of secondary metabolite (SM) backbone genes in *Aspergillus oryzae*. This figure illustrates the genomic localization of BUSCO genes and SM backbone genes in *Aspergillus oryzae*. Lines in black and green represent synteny density across 177 species and within the *Flavi* section, respectively. The gene density is shown along the chromosomes. Red bars indicate the chromosomal locations of BUSCO genes, with bar width proportional to gene length. Black and blue bars mark the positions of shared and unique SM backbone genes, respectively, with their widths corresponding to gene length. Numbers adjacent to the black bars denote SM backbone genes associated with known metabolites as listed on the MIBiG website (Table S5).

**Supplementary Figure 6.**
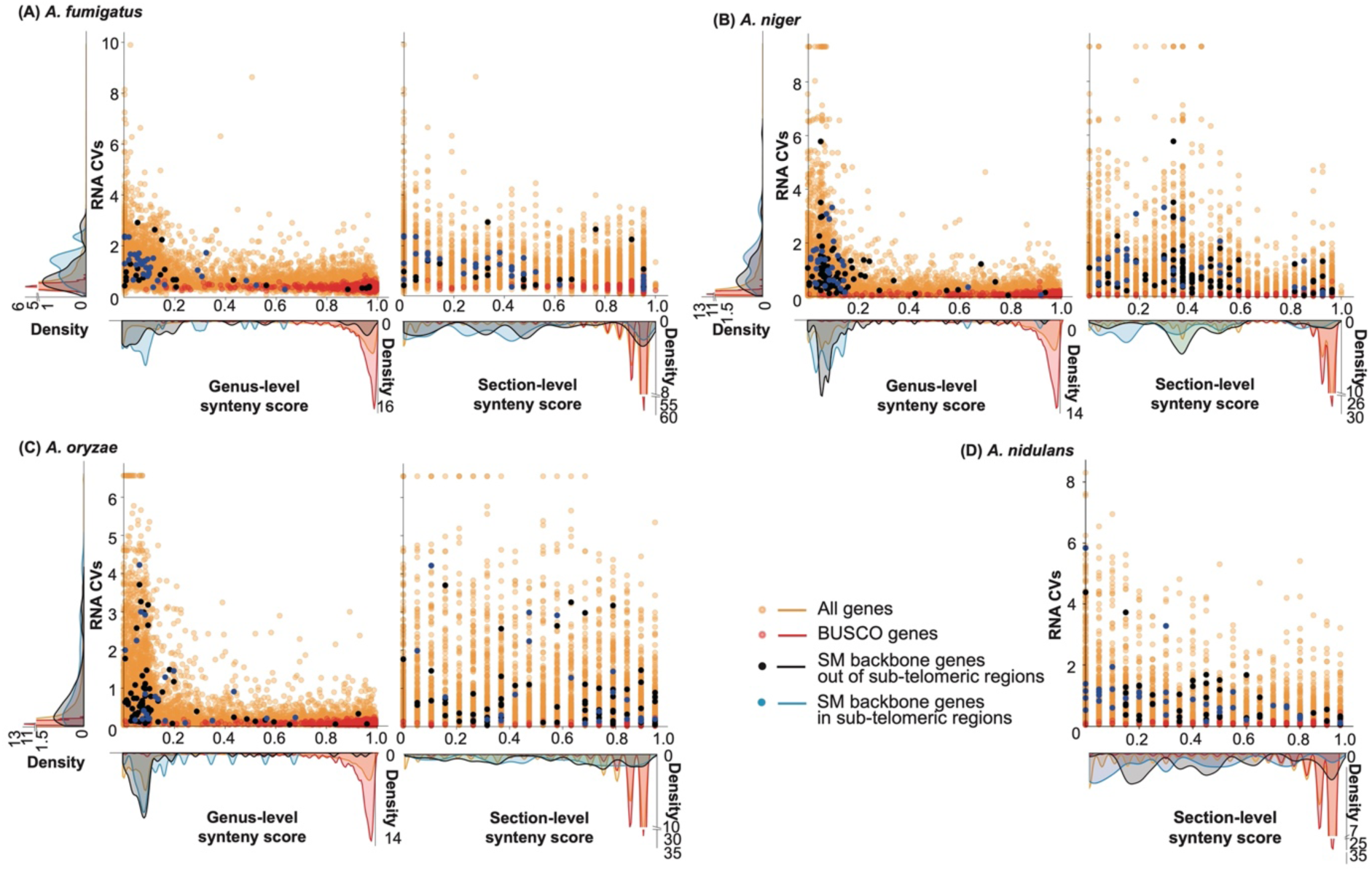
Correlation between genomic localization of secondary metabolite (SM) backbone genes and variation in gene expression in four Aspergilli. We employed the coefficient of variance (CV) to assess the relative variability of all genes, BUSCO genes, and SM backbone genes within (A) *A fumigatus*, (B) *A. niger*, (C) *A. oryzae*, and (D) *A. nidulans*. The variability is depicted using distinct colors for dots and density lines to differentiate between the gene groups. RNA-seq data was downloaded from SRA (Table S7) and ChIP-seq data is from the publication (Gacek-Matthews et al., 2016).

**Supplementary Figure 7.**
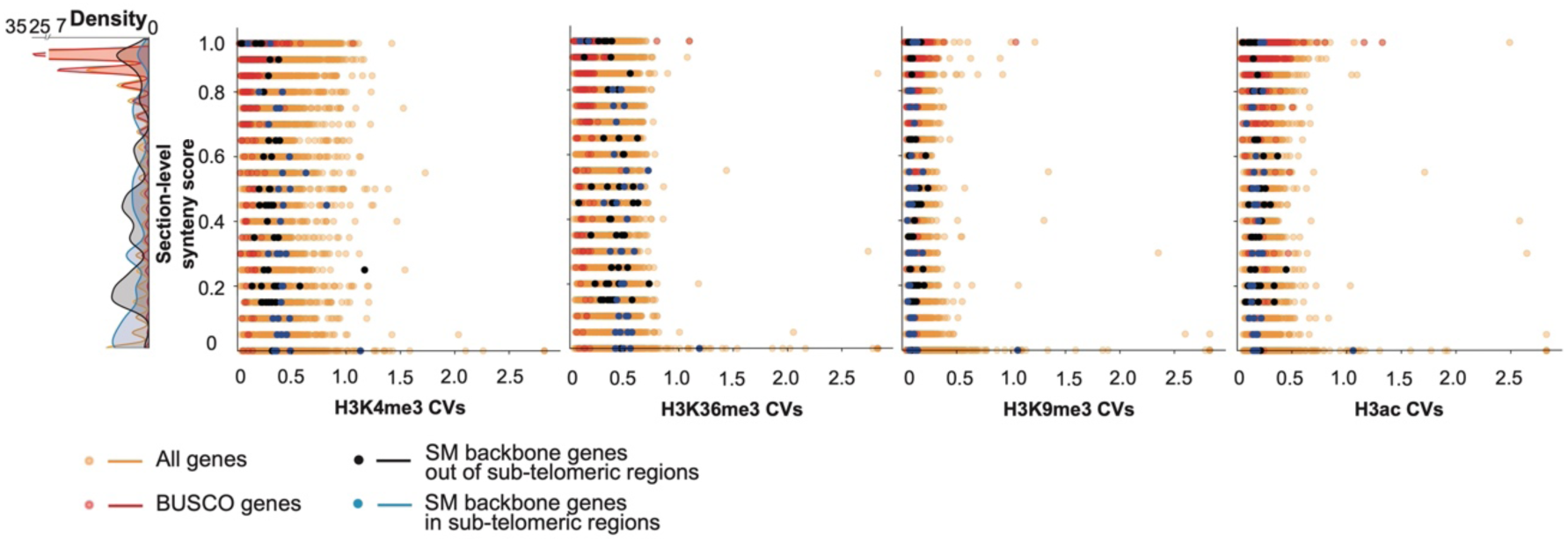
Correlation between variation in histone post-translation modifications and section-level synteny score in *Aspergillus nidulans*. We employed the coefficient of variance (CV) to assess the relative variability of all genes, BUSCO genes, and SM backbone genes in *A. nidulans*. The variability is depicted using distinct colors for dots and density lines to differentiate between the gene groups. ChIP-seq data is from the publication (Gacek-Matthews et al., 2016).

**Supplementary Figure 8.**
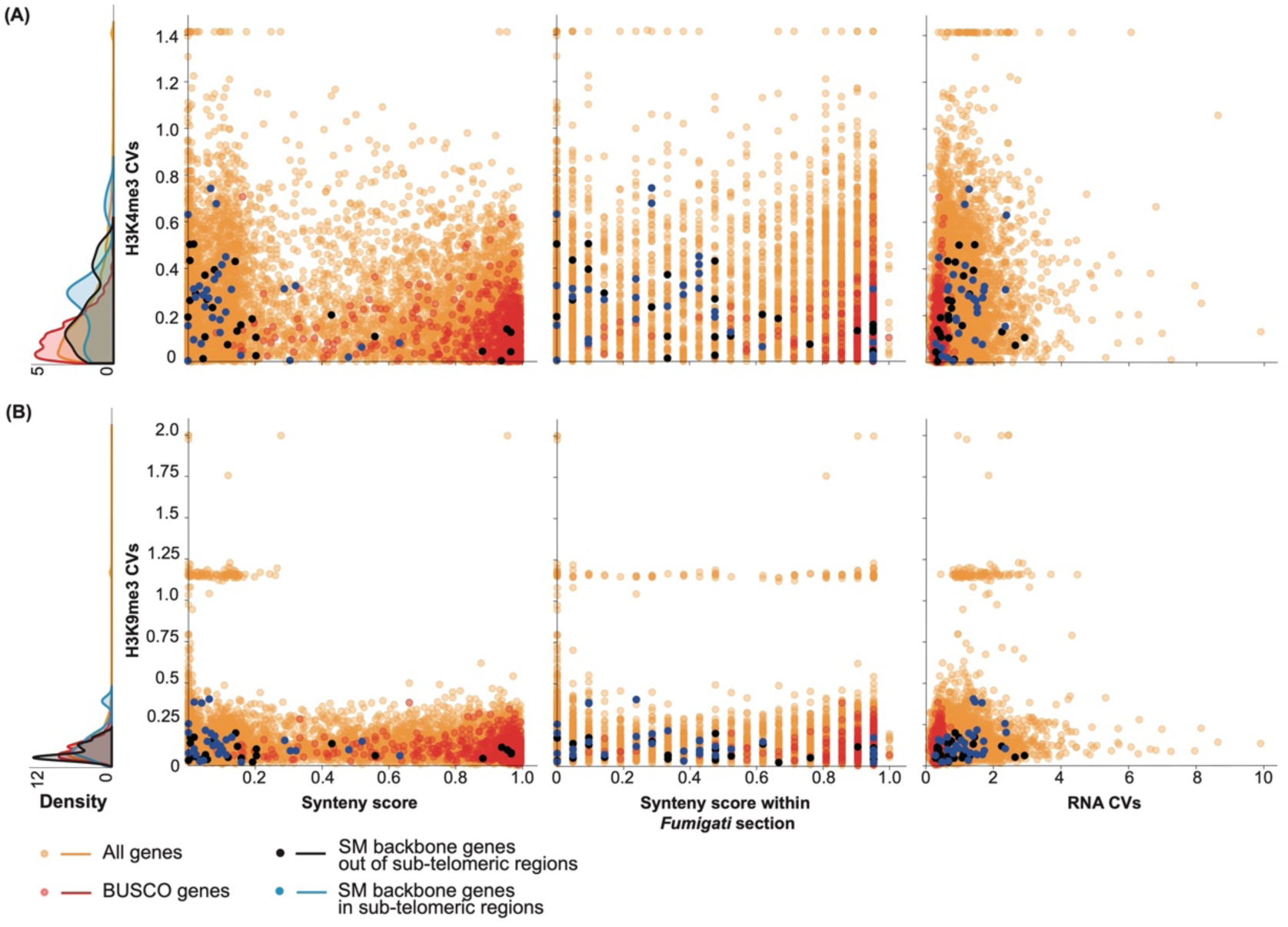
Correlation between genomic localization of secondary metabolite (SM) backbone genes and variation in gene expression and histone modifications in *Aspergillus fumigatus*. We employed the coefficient of variance (CV) of H3K4me3 and H3K9me3 to assess the relative variability of all genes, BUSCO genes, and SM backbone genes. Data is from publication (Colabardini et al., 2022). The variability is depicted using distinct colors for dots and density lines to differentiate between the gene groups.

## Data Summary

The genomes used in this study are publicly available at the JGI (Joint Genome Institute) MycoCosm repository (Grigoriev et al., 2014); the species names and abbreviations are listed in Table S1. To evaluate the completeness of the predicted proteomes and to obtain a species phylogeny, 758 fungal BUSCO (Benchmarking Universal Single-Copy Ortholog) genes were used, and their names are listed in Table S2. RNA-seq data is deposited in the SRA with the dataset identifier (RJNA1078405). Table S4 (Distribution of Orthologous groups) is deposited at Zenodo (DOI: 10.5281/zenodo.10638673).

## Acknowledgment

The sequence data were produced by the US Department of Energy Joint Genome Institute (http://www.jgi.doe.gov/) in collaboration with the user community. Especially, we thank Scott Baker, Mikael Andersen, and Joseph Spatafora for their permission to use the sequence data. We thank Jos Houbraken for validating the *Aspergillus* species tree.

## Authors and contribution

X.Z. formal analysis, investigation, methodology, visualization, writing original draft; I.L. formal analysis, investigation, methodology, visualization; J.C. conceptualization, funding acquisition, methodology, project administrations, supervision, visualization, writing original draft; M.F.S. conceptualization, funding acquisition, methodology, project administrations, supervision, visualization, writing original draft.

