## Supplementary figures and images for "Genomic localization bias of secondary metabolite gene clusters and association with histone modifications in *Aspergillus*"

### Figure S1

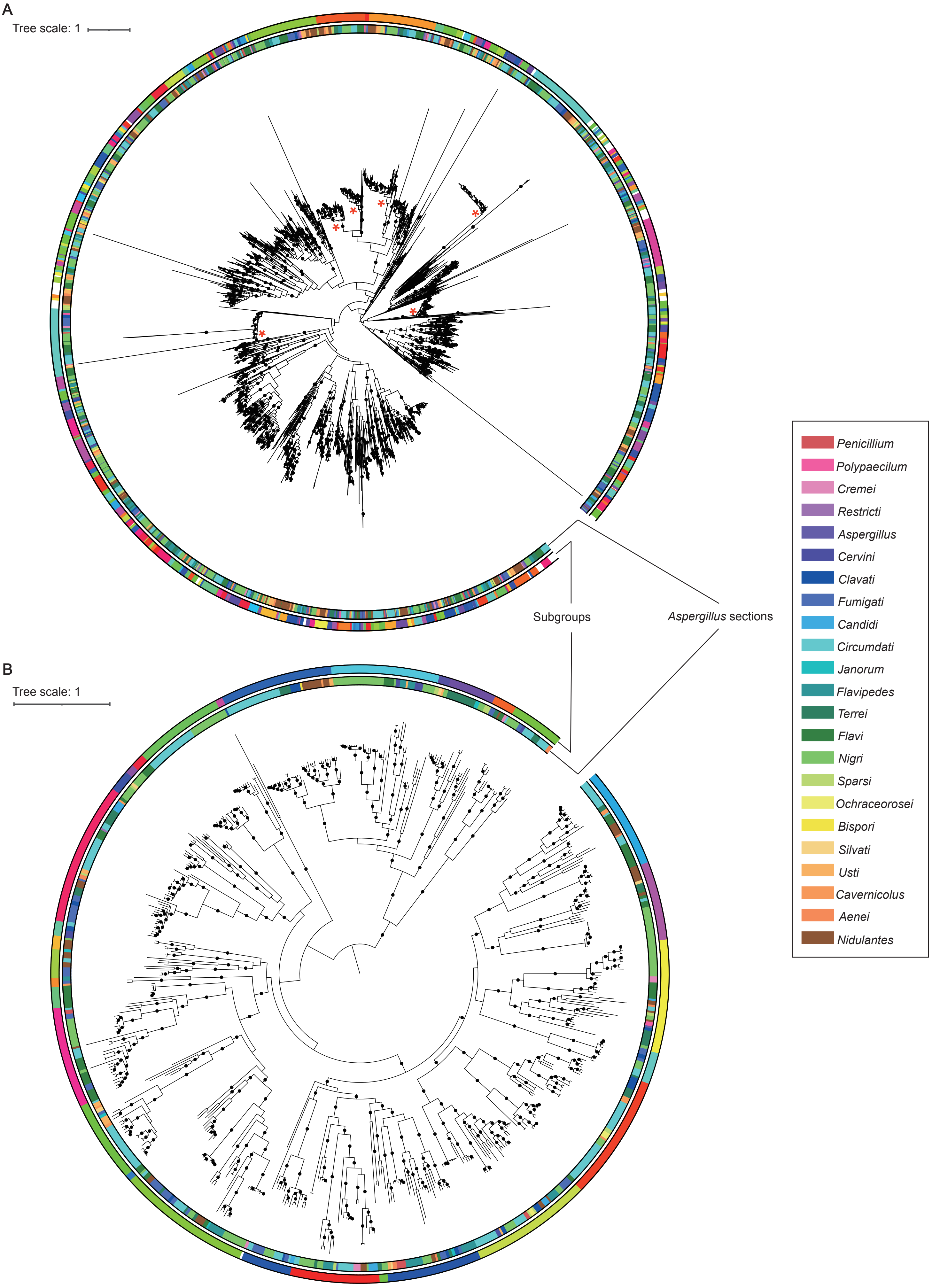


**Supplementaty Figure 1** xxx

### Figure S2

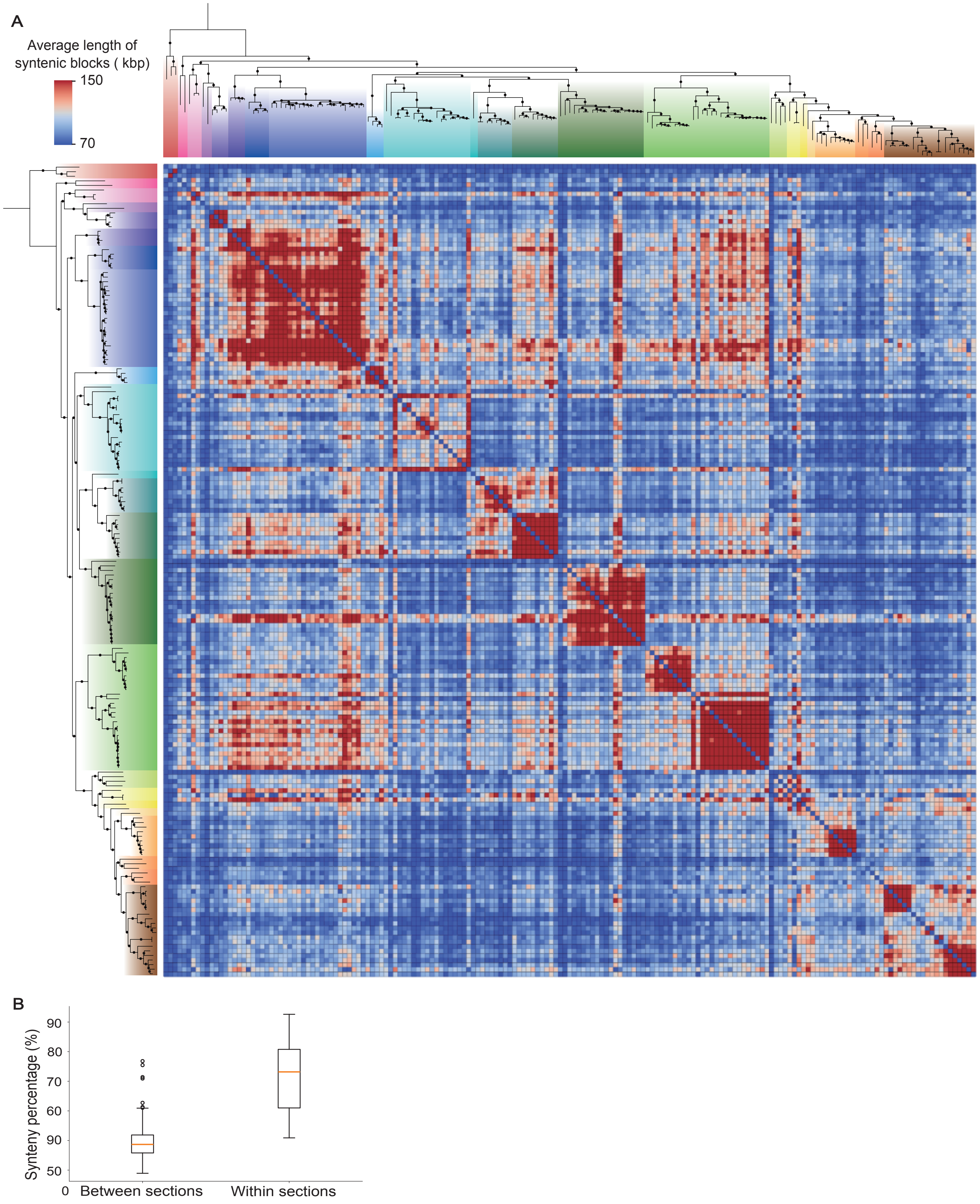

### Figure S3

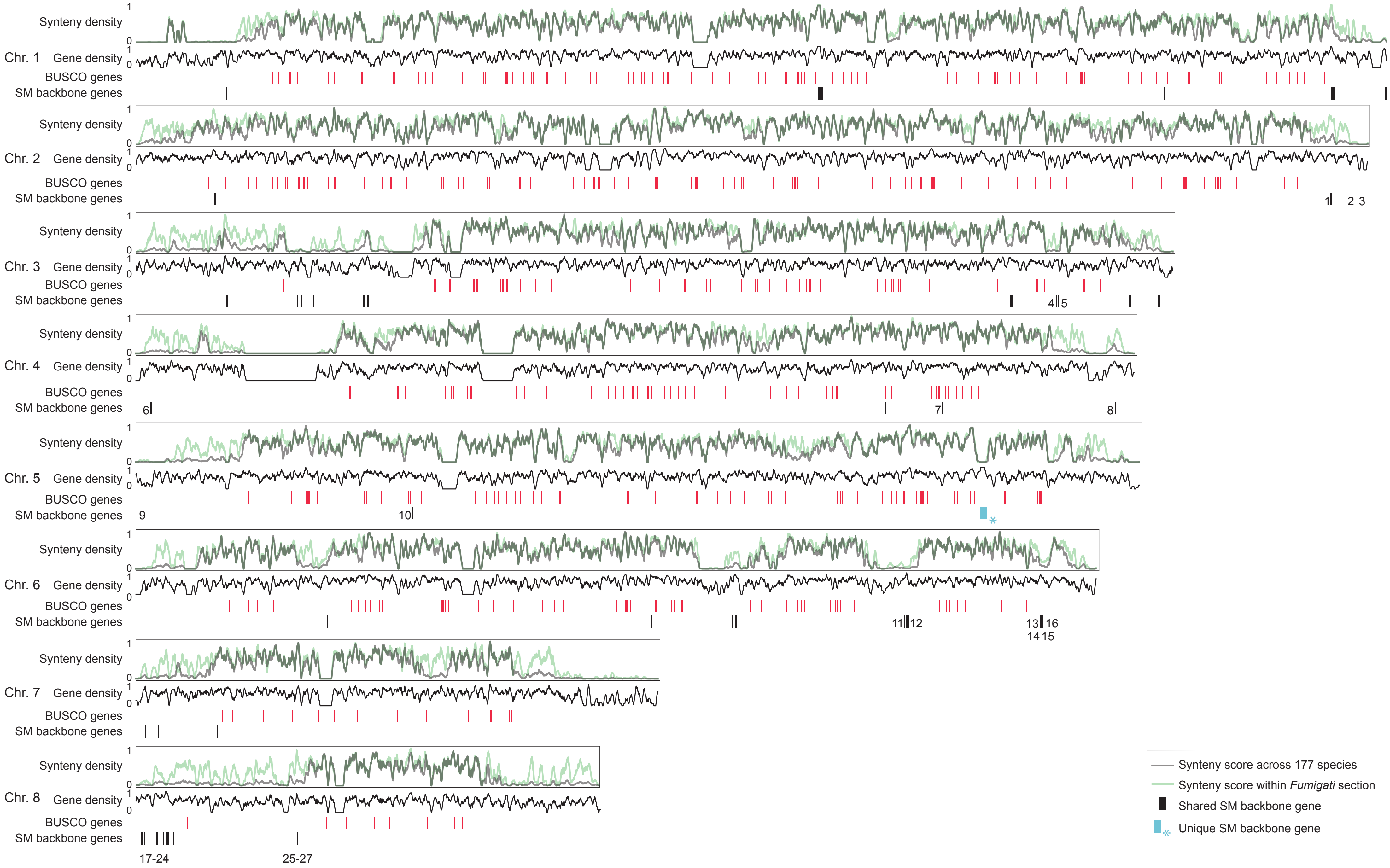

### Figure S4

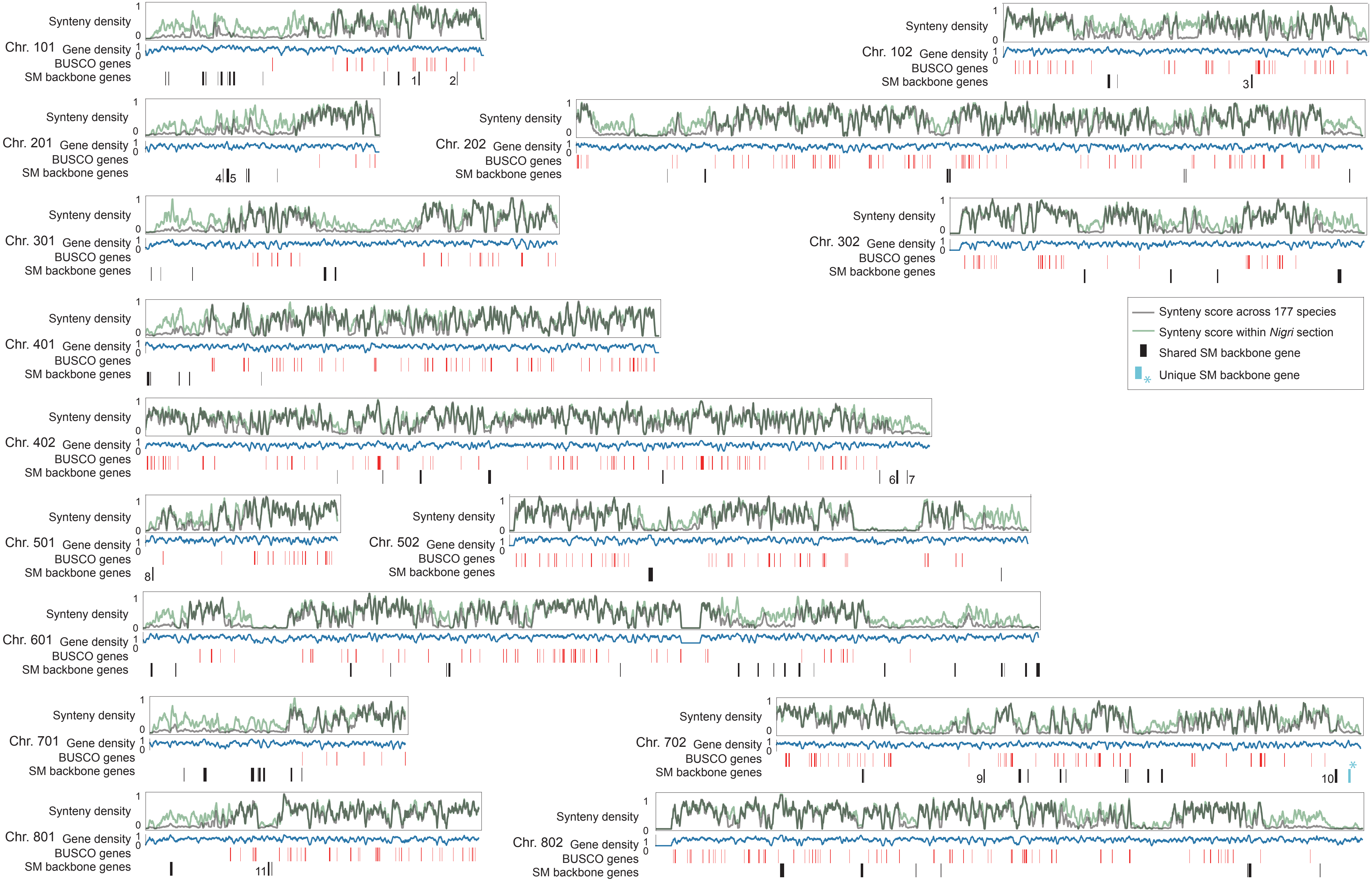

### Figure S5

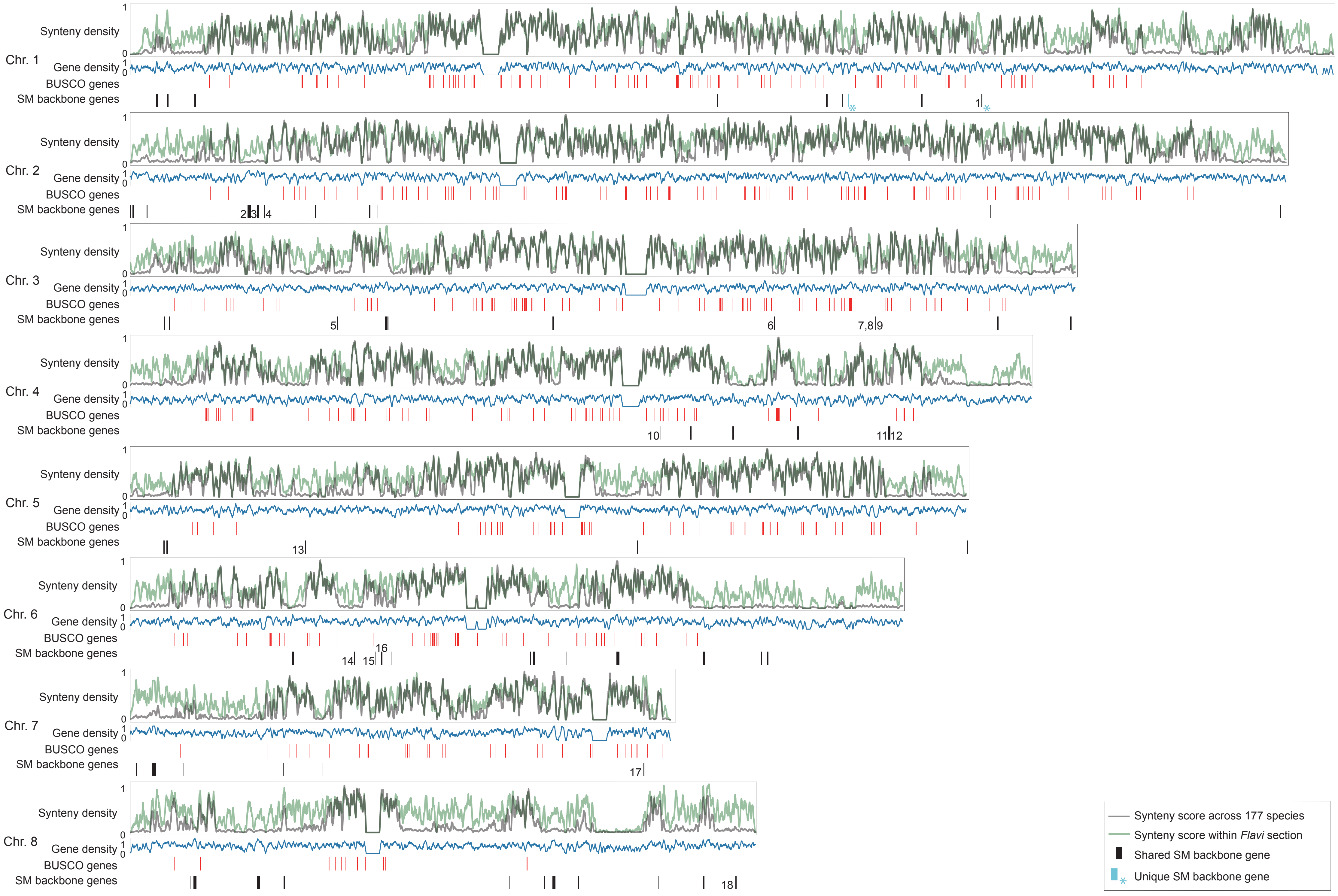

### Figure S6

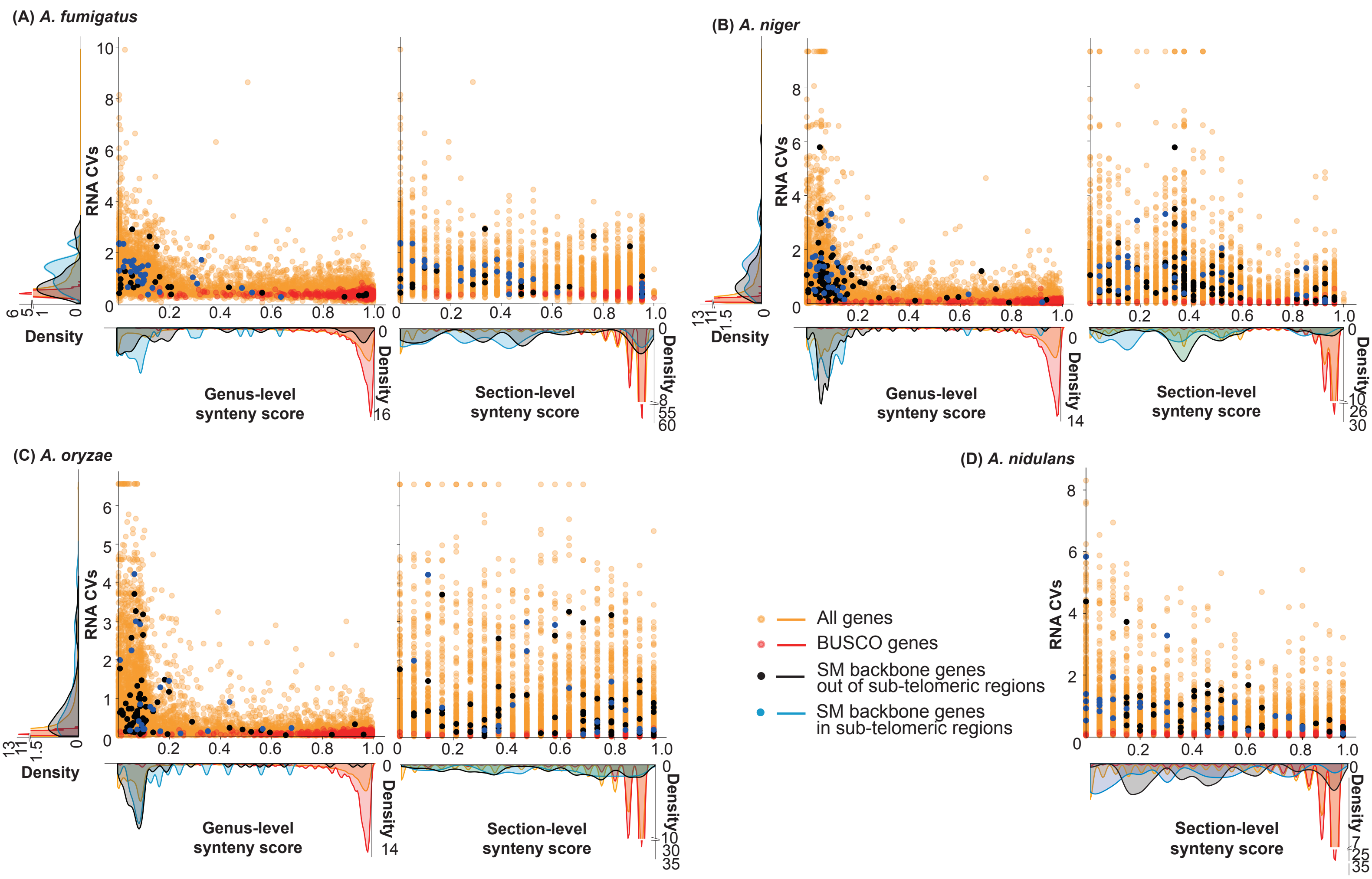

### Figure S7

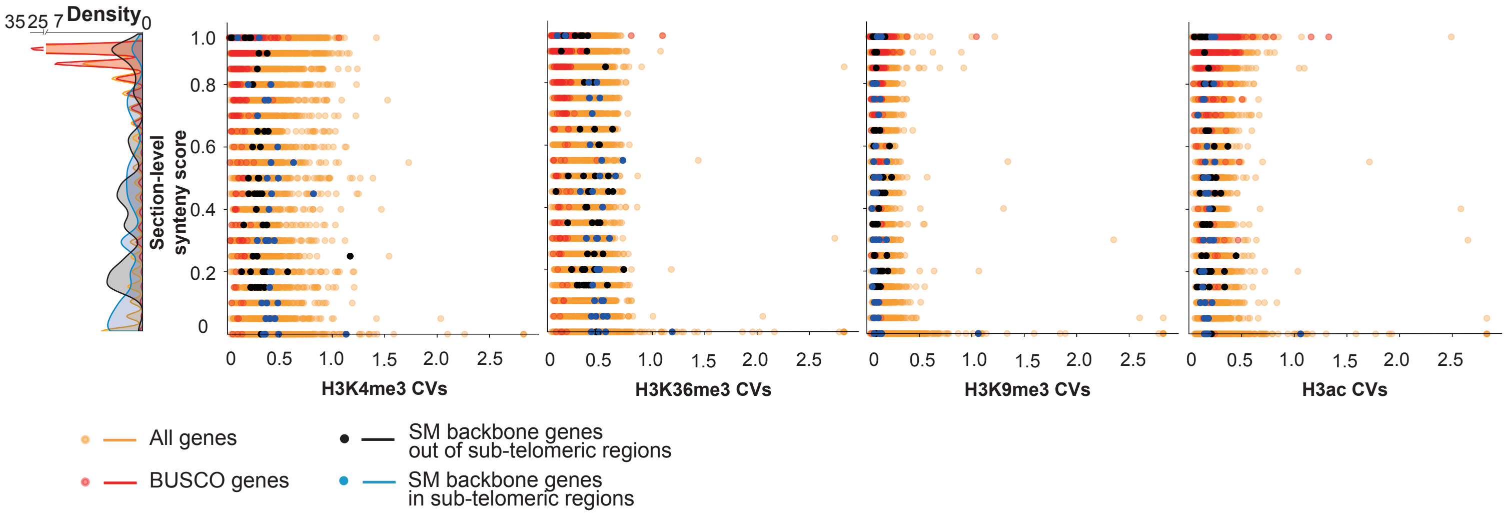

### Figure S8

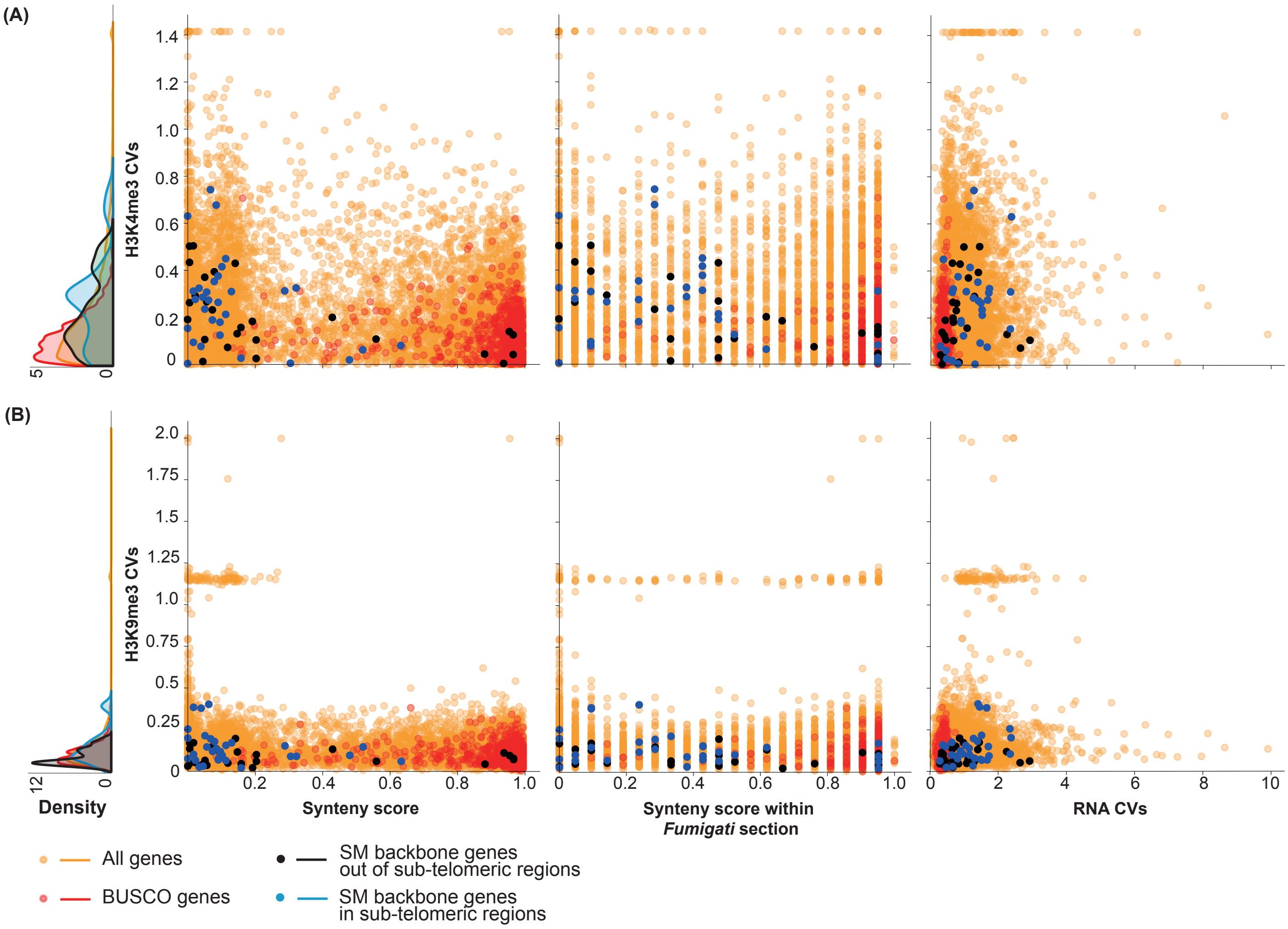
